# Large-scale analysis of the N-terminal regulatory elements of the kinase domain in plant receptor-like kinase family

**DOI:** 10.1101/2022.12.10.519927

**Authors:** Qiong Fu, Qian Liu, Rensen Zhang, Jia Chen, Hengchang Guo, Zhenhua Ming, Feng Yu, Heping Zheng

## Abstract

The N-terminal regulatory element of the kinase domain (NRE), which includes juxtamembrane segment (JM) of Receptor-like kinases (RLKs) and the N-terminal extension segment of the kinase domain in RLCKs, is the key component that regulates the activities of RLKs and RLCKs. However, their feature and functions remain largely unexplored. Herein, we perform a systematic analysis of 510,233 NRE sequences in RLKs and RLCKs from 528 plants by integrating information theory and genome-wide analysis to unravel their common characteristics and diversity. Recombinant RLKs are used to explore the structural-functional relationship of the newly-discovered motifs *in vitro*. Most NRE segments are around 40-80 amino acids, featuring a serine-rich region and a 14-amino-acid motif ‘FSYEELEKATBNFS’ which harbors a characteristic α-helix connecting to the core kinase domain. This α-helix suppresses FERONIA’s kinase activity. Motif discovery algorithm has identified 29 motifs with known phosphorylation sites highly conserved in RLK and RLCK classes, especially the motif ‘VGPWKpTGLpSGQLQKAFVTGVP’ in LRR-VI-2 class. The NRE phosphorylation of an LRR-VI-2 member MDIS1 modulates the auto-phosphorylation of its co-receptor MIK1, indicating NRE’s potential role as a “kinase switch” in RLK activation. Consequently, the characterization of phosphorylatable NRE motifs improves the accuracy in predicting phosphorylatable serines. Altogether, our study provides an enriched and comprehensive dataset to investigate NRE segments from individual RLKs and helps understand the underlying mechanism of action of the RLK signal transduction and kinase activation processes in plant adaptation.

## Introduction

Receptor-like kinases (RLKs) superfamily is the largest class of plant kinases responsible for signal transduction across the plasma membrane. RLKs regulate a wide array of signaling pathways that involve plant growth and development, perception of external signals, and response to adverse environments. The earliest description of RLK dates back to the 1990 study describing a set of kinases from maize with similar characteristics as the receptor tyrosine kinase (RTK) from animals (Walker and Zhang, 1990). The RLK discovered in maize exhibits typical RLK features containing a larger extracellular domain (ECD) and transmembrane helix on top of the cytoplasmic kinase domain. However, some plant RLKs lack apparent extracellular domain and/or transmembrane regions and are referred to as Receptor-like Cytoplasmic Kinases (RLCKs) (Dievart et al., 2020). A systematic study based on the kinases domain phylogeny in the model organism *Arabidopsis thaliana* indicates that RLKs and RLCKs are a large family of plant kinases (Shiu and Bleecker, 2001). A study of Expression Sequence Tag (EST) from various land plants indicates that the RLK gene superfamily emerged early on the evolutionary path and started expansion before the diversification of land plants from their aquatic ancestors (Gong and Han, 2021).

Further research reveals that members of the RLK superfamily in plants share similar features with the gene product of *Pelle* from *Drosophila melanogaster* and the Interleukin Receptor-associated Kinases (IRAKs) from mammals (Belvin and Anderson, 1996; Cao et al., 1996). Although *Pelle* and IRAKs in animals are much less common than their RLK counterparts in plants, their similarity suggests that they most likely have a common ancestor from the evolutionary perspective (Shiu and Bleecker, 2001). The plant RLK superfamily, mammal IRAKs, and *Pelle* from *Drosophila melanogaster* form the RLK/Pelle monophyletic clade of kinases as cataloged by the Shiu protein kinase classification scheme (Shiu and Bleecker, 2003). This scheme subdivides the plant RLK superfamily into 65 classes, of which the most commonly observed features are RLCKs (20 classes) and Leucine-rich repeat (LRR) kinases (23 classes). The RLK superfamily has undergone highly complex evolutionary changes due to extensive gene expansions coupled with functional divergence in response to various environmental signals (Lehti-Shiu and Shiu, 2012).

Structural topology and domain organization of RLKs typically encompass an extracellular domain mainly responsible for sensing various external signals, a single transmembrane region, and a cytoplasmic kinase for activating downstream events. RLKs are versatile in sensing a wide array of external events while encountering partner proteins, peptides, carbohydrates, organic small molecule agents, and inorganic chemical or physical signals. According to the extracellular domain, the most prominent RLK classes are S-domain, LRR, and Epidermal growth factor-like repeat class (Walker, 1994). The S-domain class is characterized in Brassicaceae during the study of self-incompatibility-locus glycoproteins (SLGs), a critical component in preventing self-pollination (Nasrallah and Nasrallah, 1993). LRR kinases play various roles in plant signal transduction processes, benefiting from the tandem repeats of highly conserved β-sheets facilitating extensive protein-protein interactions (Kobe and Deisenhofer, 1993). Furthermore, RLCKs can be associated with RLKs to regulate biotic and abiotic processes (Liang and Zhou, 2018). In particular, RLCKs can act as common signaling nodes that link RLKs to downstream signal output. Recent studies have suggested a phosphorylation relay in which RLKs phosphorylate RLCKs upon ligand perception, and the RLCKs further phosphorylate major downstream signaling components (Du et al., 2016; Li et al., 2017).

The intrinsically disordered region between the transmembrane region and the cytoplasmic kinase in members of the RLK superfamily is called the juxtamembrane (JM) segment. The structurally flexible JM critically influences cytoplasmic conformational changes and is highly sensitive and responsive to phosphorylation, activating the downstream kinase activities. In the epidermal growth factor (ErbB-1) kinase, deleting the JM segments causes a severe loss of tyrosine phosphorylation (Thiel and Carpenter, 2007). In the Eph receptor kinase (EphB2), the two conserved tyrosines (Y605 and Y611) in the JM segment regulate the kinase autophosphorylation (Binns et al., 2000; Zisch et al., 2000). In type I transforming growth factor (TGF-β) receptor kinase, phosphorylation of the JM segments prevents the binding of PKBP12 and activates the kinase activity of TGF-β (Hubbard, 2001; Huse et al., 2001). In *Arabidopsis* and rice, the JM segments of CERK1 and other LysM-RLKs are heavily involved in the chitin-induced signaling process (Zhou et al., 2020), while the phosphorylation of this JM segment activates the mechanism to protect the plant against microbial infection (Gong et al., 2019). The OsWAK11-mediated inhibitory phosphorylation occurs within an STS motif (Ser751-Thr752-Ser753) localized to the JM of OsBRI1 found in most monocots (Yue et al., 2022). While many RLCKs lack a transmembrane region and hence do not possess a JM segment by definition, the vast majority of RLCKs still exhibit a characteristic disordered region as an extended sequence in the N-terminus of the kinase domain (NKE). NKEs share many typical regulatory roles as the JM segments in RLKs, as PTMs and conformational changes of the NKE may also activate or inhibit the RLCK activities. For example, the NKE region of BIK1 protein includes ten phosphorylation sites that could be phosphorylated by SIK1 to increase BIK1’s stability (Zhang et al., 2018). In this study, JM sequences and the N-terminal extension sequences are collectively referred to as putative N-terminal regulatory elements of the kinase domain (NRE).

It becomes increasingly apparent that the NRE is a crucial component correlating kinase activation with phosphorylation and other posttranslational modification (PTM) of cytoplasmic regions in RLKs and RLCKs. Despite the versatile and critical functions of NRE segment, there have been few systematic studies of the NRE. The intrinsically disordered nature of the NRE segment adds another level of complexity to the problem due to its conformational flexibility. Herein, we systematically investigate NRE segments’ occurrence and sequence features from various aspects, aiming to characterize and annotate NRE sequences in the plant RLK superfamily. Our data mining of the genomic and structural data available for NRE segments in the RLKs superfamily provides a dataset that facilitates further investigation of NRE segments from individual RLKs and RLCKs. An in-depth understanding of the NREs can help understand the underlying mechanism of action and the structural-functional relationship of the RLK and RLCK kinase activation alongside the signal transduction process in plants.

## Results

### Occurrence of putative N-terminal regulatory elements of the kinase domain in plant RLK superfamily

The occurrence of NRE, which includes both JM and NKE, is investigated in various plant groups and all RLK and RLCK classes (Fig. 1) as cataloged by the Shiu scheme (Shiu and Bleecker, 2001). Each member from the RLK superfamily is annotated with one of the three mutually exclusive topological groups according to the presence or absence of the transmembrane region and the length of ECD: RLK, RLCK with TM region, RLCK without TM region (RLCK with NKE). Our non-redundant RLK and RLCK dataset identified from the genomic sequence of 528 plant species results in the presence of 313,468 RLKs, 46,375 RLCK with TM region, and 150,390 RLCK without TM region (Fig. 1). RLCK without N-terminal extension (NKE) is rarely observed with only <2000 instances and is excluded from further analysis in this study. In total, our dataset has 359,843 JM sequences and 150,390 NKE sequences.

**Figure 1.**
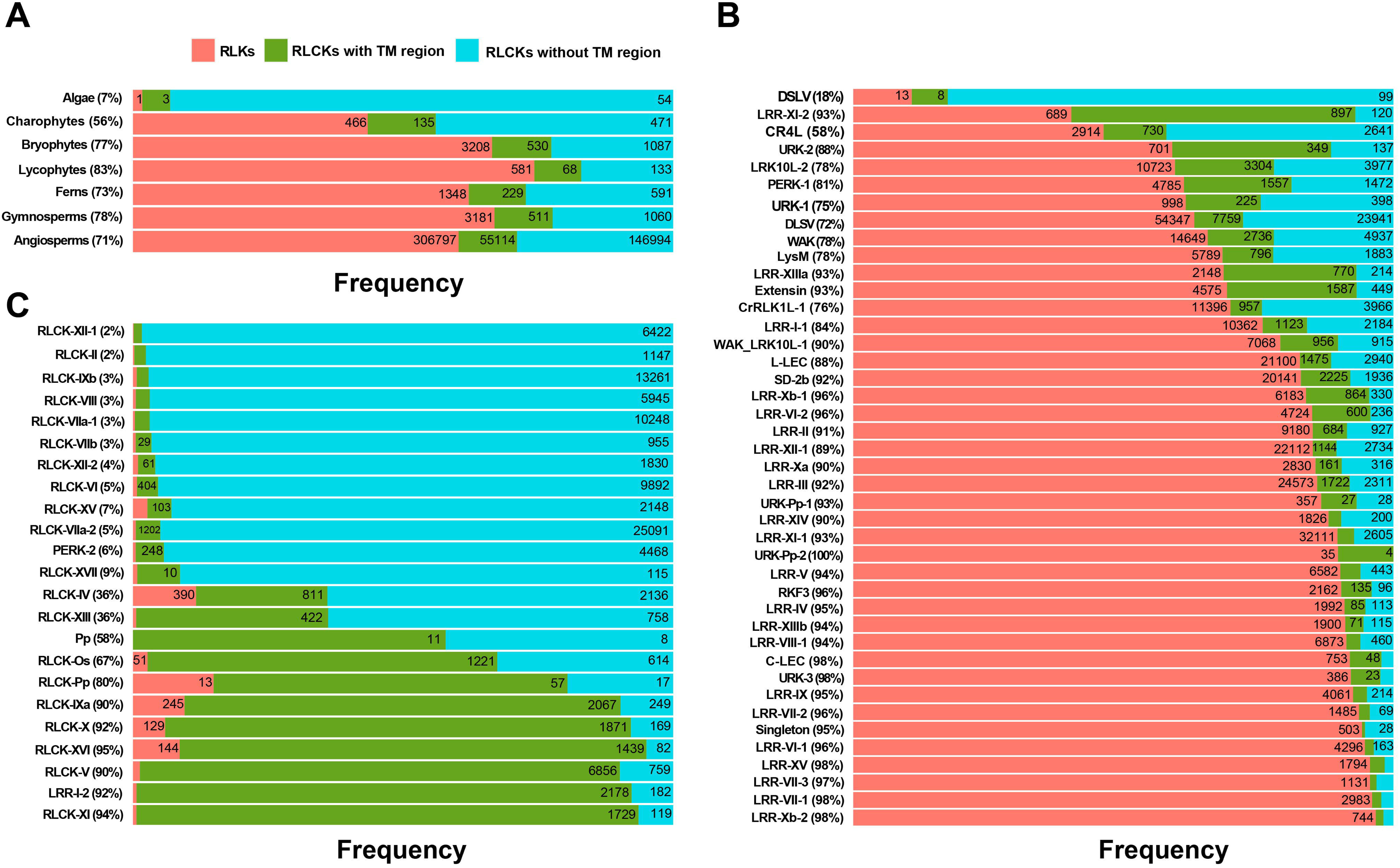
The occurrence of NRE segments in the RLK superfamily. (**A**) The emergence of NRE segments in different plant groups. (**B**) The emergence of NRE segments in different classes of RLKs. (**C**) The emergence of NRE segments in different classes of RLCKs. The classes used the designation the kinase classification scheme proposed by Lehti-Shiu and Shiu. Red box refers to RLK sequences with extracellular domains (ECD), transmembrane regions (TM), and intracellular domain (CD). Green box refers to RLCK sequences with limited length of ECD and with TM and CD. Cyan box refers to RLCK sequences without ECD and TM. Percentage refers to the proportion of JM.

In algae, RLCKs having NKE segment is the main component, as evidenced by 54 RLCKs without TM region of a total of 58 RLKs and RLCKs (Fig. 1A). Charophytes, Bryophytes, Lycophytes, Ferns, Gymnosperms, and Angiosperms exhibit abundant RLK sequences, with a lower proportion of RLCK without TM region, and an even lower proportion of RLCK with TM region. The enrichment of RLK sequences could be mainly due to the accelerated gene duplications coupled with diversifications of RLK classes compared to RLCK classes during the evolution to develop the characteristic vascular bundle and seeds features in response to the external environments (Zhang et al., 2020b).

The abundance of NRE segments is analyzed in different classes of RLKs and RLCKs according to the Shiu classification scheme (Lehti-Shiu and Shiu, 2012) (Fig. 1B, C). The RLK classes are mainly characterized by a high percentage (72%-100%) of RLKs or RLCKs with TM region, which defines the presence of a JM segment, except for DSLV, LRR-XI-2, and CR4L (Fig. 1B). Most of the RLCK classes lack ECD, yet may exhibit either the absence of TM region or the presence of TM region. For example, RLCK classes with more than 50% of the sequences showing a TM region include Pp, RLCK-Os, RLCK-Pp, RLCK-IXa, RLCK-X, RLCK-XVI, RLCK-V and RLCK-XI (Fig. 1C). Besides all RLCK classes, a few RLK classes are also assigned into the topology of RLCK, including PERK-2, Pp, and LRR-I-2 (Fig. 1C). While many RLKs that belong to either the PERK-2 class or the Pp class lack a characteristic TM region, more than 90% members of the LRR-I-2 class do possess a TM region yet exhibit an exceptionally short ECD sequences not qualified to be considered as an ECD domain by our topological assignment algorithm. Further investigation indicates that the vast majority (about 72.2%) of the LRR-I-2 class members do not possess any LRR motifs. The assignment of this class as an LRR class could be due to the insufficient data availability of the original RLK classification study (Lehti-Shiu and Shiu, 2012).

### Length and composition of NRE segments in RLKs and RLCKs

Analysis of the length of NRE segments indicates that most NRE sequences of RLKs and RLCKs with TM region have a length between 30-100 amino acid residues, with the most typical length at around 40 amino acids and a secondary peak at about 50 amino acids, yet with a decreasing abundance when sequence length increases (Fig. 2A, B). In RLCKs with TM region, other lengths peaks around 80 and 140 amino acids are also observed. The length of NRE segments in RLCKs without TM region does not exhibit any noticeable peak between 14 and 100 amino acids (Fig. 2C). The distribution of NRE sequence length could vary significantly between different classes of RLKs and RLCKs defined in the Shiu classification scheme. In most classes with the largest numbers of RLK and RLCK, the most typical NRE sequence length is around 50 amino acids (Fig. S1). Some RLK and RLCK classes do exhibit a flattened distribution of NRE sequence length in the range of 60-200 amino acids, such as LRR-V, RLCK-VI, and RLCK-IXb.

**Figure 2.**
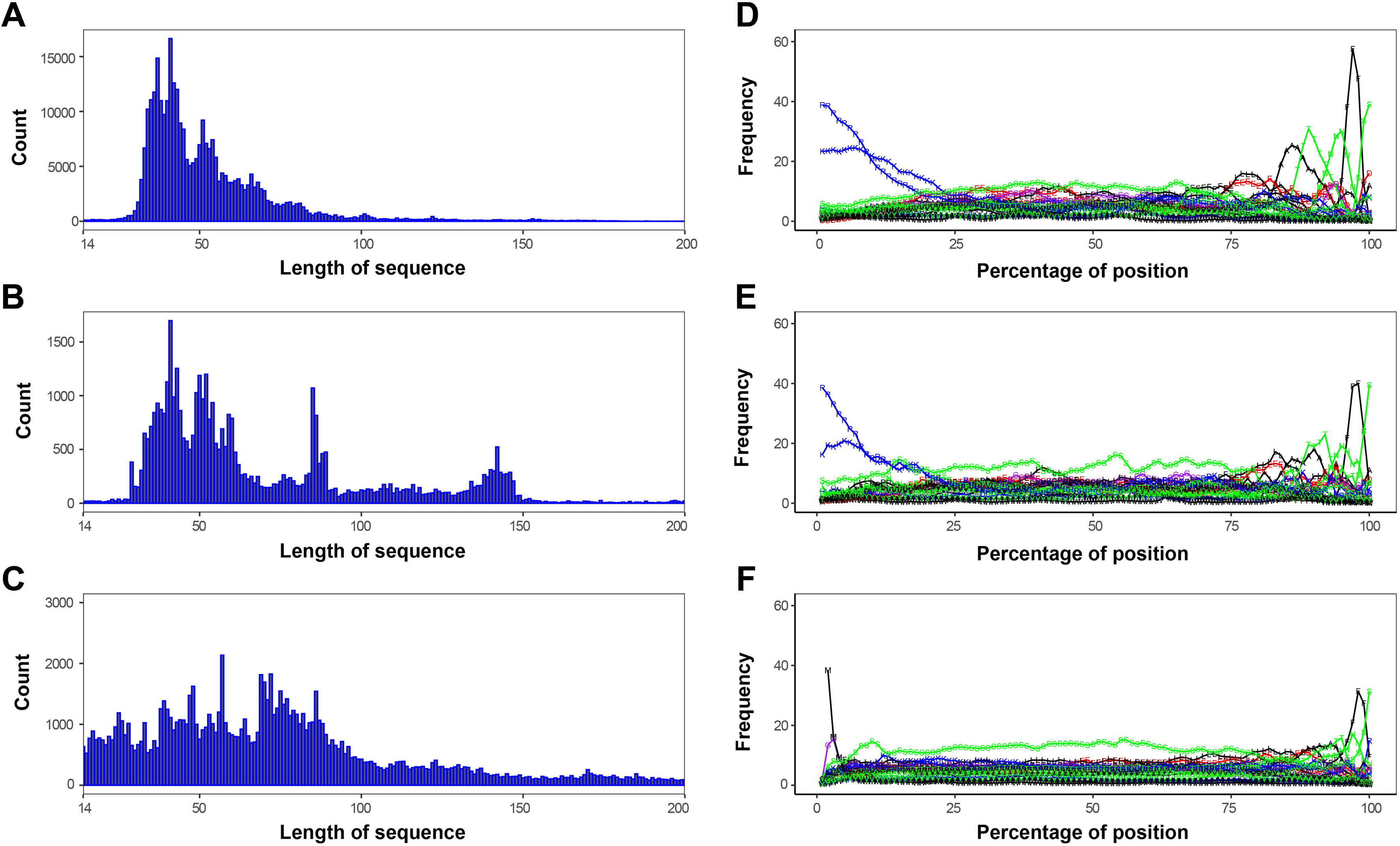
Length distribution and composition of NRE segments in the RLKs superfamily. (**A-C**) Length distribution of NRE segments. (**D-F**) Composition of NRE segments. (**A, D**) represent RLK sequences. (**B, E**) represent RLCKs sequences with TM region. (**C, F**) RLCKs sequences without TM region. The various lengths of different NRE segments are scaled to 200 to align both N-terminal and C-terminal. The 1-letter-code of the amino acid with lines connected along the NRE segment indicates the composition percentage at a specific position. Hydrophobic, polar, acidic, basic, and neutral residues are shown in black, green, red, blue, and purple, respectively.

The composition of different types of amino acids is plotted along the NRE sequence (Fig. 2D-F). The composition of RLKs and RLCKs with TM region is the same, and their N-terminal sequences are overwhelmed with almost 60% in basic residues arginine (R) and lysine (K). This trend gradually decreases to ∼10% arginine and ∼10% lysine for the first 20% along the entire length of the NRE sequence. The N-terminal composition of RLCKs without TM region is enriched in methionine (M) which is the first residue in translation. However, their C-terminal sequences exhibit identical several characteristic residues with a single peak above 20% abundance in composition: alanine (A), threonine (T), asparagine (N), phenylalanine (F), and serine (S), sequentially. The observation of multiple peaks indicates the presence of a common C-terminal sequence motif for all NRE segments for RLKs and RLCKs sequences. The whole NRE sequence boasts a consistently high serine (S) composition, with a broad plateau above 10% between 20%-80% along the NRE sequence, which may indicate the versatile role of NRE as a substrate that is subject to serine phosphorylation. Other amino acids that exhibit peaks at above 10% in the composition include glutamic acid (E) and leucine (L) - both exhibiting a primary peak at about 80% along the NRE sequence and a secondary peak at about 30%-40% along the NRE sequence.

### Sequence features of NRE segments from different classes of RLKs and RLCKs

WebLogo analysis of the amino acid profiles of the NRE sequence reveals the presence of high ‘RK’ content at the N-terminal in RLK and RLCK with TM region and the presence of high ‘M’ content at the N-terminal in RLCK without TM region, and the presence of a 14-amino acid motif “FSYEELEKATBNFS” at the C-terminal for all RLKs and RLCKs sequences, where ‘B’ refers to aspartic acid (D) or asparagine (N) (Fig. 3), in agreement with the composition analysis (Fig. 2D-F). Besides the arginine, lysine, and methionine at the N-terminal, a characteristic neutral residue glutamine (Q) / asparagine (N) is also observed in some RLK and RLCKs classes defined in the Shiu classification scheme (Fig. S2). For example, Q is observed at the N-terminal of classes LRR-I-2, LRR-IV, LRR-VIII-1, LRR-XIIIb, RLCK-VIIa-2, URK-3, and URK-Pp-2, while N is observed at the N-terminal of classes LRR-I-2, LRR-VI-2, LRR-VII-1, LRR-XIIIb, and Pp.

**Figure 3.**
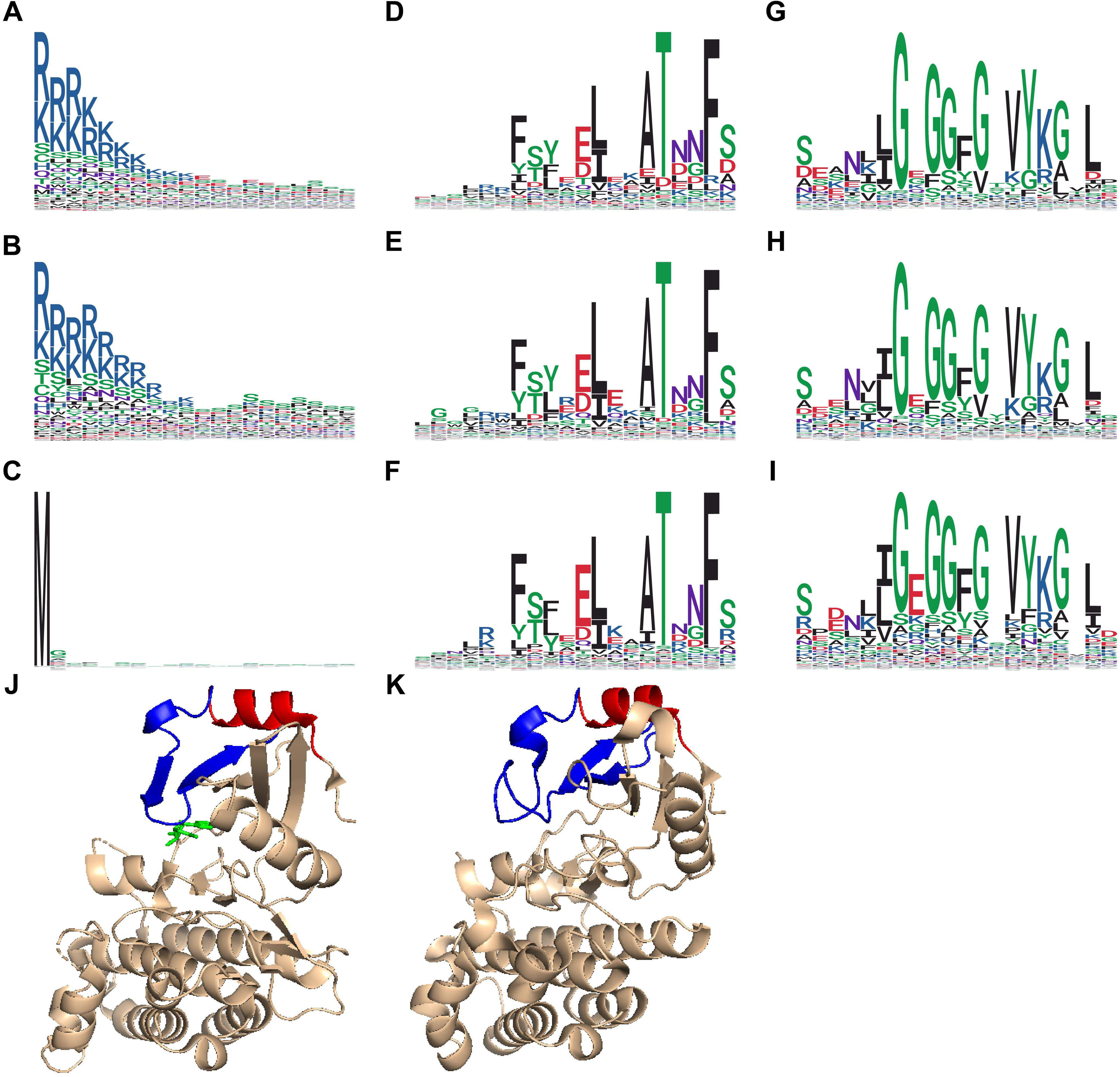
Characteristic sequences of NRE segments in the RLKs superfamily. (**A-C**) Characteristic sequences at the N-terminal of NRE segments. (**D-F**) Characteristic sequences at C-terminal of NRE segments. (**G-I**) Characteristic sequences at the N-terminal of the kinase domain in RLK. (**A, D, G**) represent RLK sequences. (**B, E, H**) represent RLCK sequences with TM region. (**C, F, I**) represent RLCK sequences without TM region. (**J, K**) Structure of the kinase domain with partial NRE segment (C-terminal) of RLK and RLCK, with C-terminal of the NRE segment colored in red, and N-terminal of the kinase domain colored in blue. Hydrophobic, polar, acidic, basic, and neutral residues are shown in black, green, red, blue, and purple, respectively.

The C-terminal of the NRE sequence exhibits a conserved 14-amino acid motif “FSYEELEKATBNFS” immediately adjacent to the N-terminal of the protein kinase domain. The threonine at −5 position is conserved among all RLK and RLCK NRE segments, except for LRR-III, LRR-VI-2, LRR-XI-2, LRR-VII-1, RLCK-II, and Singleton classes (Fig. S3). In these classes, threonine −5 is primarily substituted by leucine, cysteine, aspartic acid, and phenylalanine.

A single serine residue is found at the C-terminal −1 position of the NRE segment as the last residue of the motifs. While in the middle of the NRE segment, serine-rich regions are found up to 25% abundance beginning at 20% of the NRE sequence and ending at 60% of the NRE sequence, especially in classes C-LEC, DSLV, Extensin, LRR-I-2, LRR-IV, LRR-IX, LRR-VII-1, LRR-VIII-1, LRR-Xa, LRR-Xb-2, LRR-XIIIa, LRR-XIV, RLCK-IV, RLCK-V, RLCK-XVI, Singleton, URK-1and URK-Pp-2 (Fig. S3). A single leucine-rich region is found close to the C-terminal of the NRE segments in LRR-III, LRR-VII-2, LRR-Xa, LRR-Xb-1, LRR-XIIIa, RLCK-IXa. An additional leucine-rich region is also found in the middle of the NRE segments in LRR-II, LRR-VI-1/2, LRR-XIIIb, WAK (Fig. S4).

### Characteristic α-helix of the NRE segment modulates FER kinase activity

A typical and well-studied RLK FER is selected to evaluate the relevance of the universal α-helix of the NRE segment in RLK. Two constructs are prepared to recombinantly express the kinase domain of FER, either with the α-helix (residues 518-816) or without the α-helix (core kinase, residues 536-808) (Fig. 4A). FER K565R mutant does not have kinase activity and is thus used as a negative control. The FER^KD^ core kinase (residues 536-808) consumes ATP faster than FER with the α-helix of the NRE segment (residues 518-816) (Fig. 4B). Our data suggest that the α-helix of the NRE segment may negatively regulate FER’s kinase activity via suppressing its ATPase activity.

**Figure 4.**
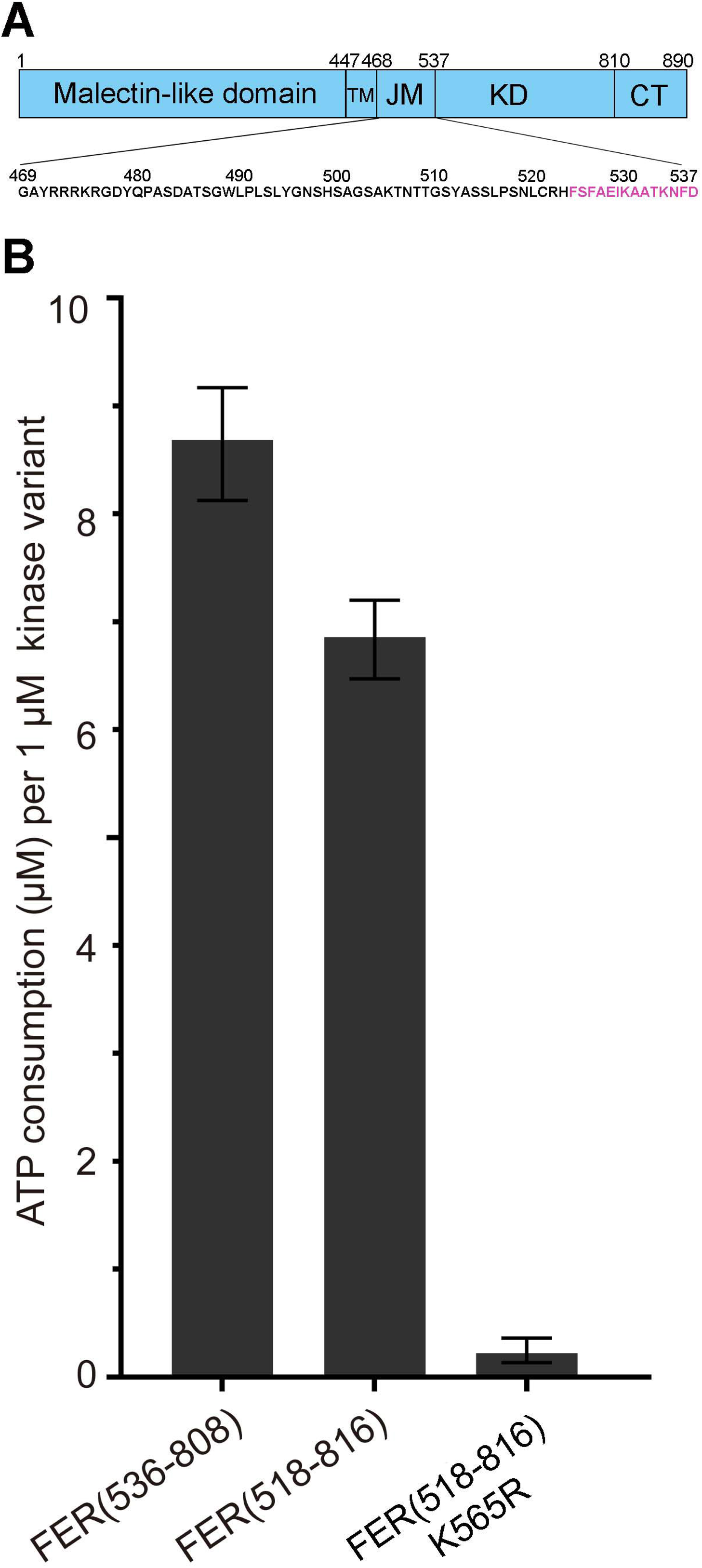
Characteristic α-helix of the NRE segment modulates FER kinase activity. (**A**) Schematic representation of FER RLK. The amino acids marked in purple represent the C-terminal α-helix of the NRE segment. (**B**) The kinase activity of the FER constructs.

### Characteristic motifs in the NRE segments

MEME identifies 65 motifs in the NRE segments with a *p*-value threshold of 0.005 and a normalized frequency higher than 1. All identified motifs that typically exist in modern embryophytes are found in early diverging streptophyte algae (Charophytes), while only 14 of the 65 motifs are also identified in algae (Fig. S5). Out of 65 analyzed, 31 motifs have experimentally determined phosphorylation sites associated with abiotic stress responses, development, and plant-pathogen interactions in embryophytes (Table S3). Meanwhile, these motifs from algae refer mainly to cellular energy signaling, plant defense signal transduction, and environmental stimuli response (Meyer et al., 2012; Lv et al., 2014; Roitinger et al., 2015; Nukarinen et al., 2016; Yuan et al., 2016). Among these motifs, 21 motifs are of single (39%) or double (29%) phosphorylation sites; however, ten motifs have 3-9 phosphorylation sites (32%). The low frequency of tyrosine phosphorylation is likely attributed to the reports of its frequent (75%) multiphosphorylation (Sugiyama et al., 2008). Indeed, our analysis reveals the presence of only phosphotyrosine sites in four multiphosphorylated motifs of all 31 motifs analyzed (Table S3).

Sequence similarity between the phosphorylation motifs and the matching NRE sequences is used as an additional criterion to screen the most conserved phosphorylation motifs for further functional analysis. Out of 31 motifs with experimentally determined phosphorylation sites, 29 motifs show up to 50% similarity with the matching NRE sequences (Table 1). The top four motifs with the highest scores for the mean sequence similarity (SI) include one motif (‘VGPWKpTGLpSGQLQKAFVTGVP’) observed in LRR-VI-2 class, one motif (‘KEPLpSINVATFEKPL’) observed in LRR-Xb-1 class, and two motifs (‘pTSSEQKSDITDpSCSQMILQLHDVYDPNKINVKIKIVSGSPC’ and ‘SIRGPVVpTPTpSpSPELGTPFTATEAGTSSVSSSDPGTSPFFI’) observed in PERK-2 class. The highest-scored motif ‘VGPWKpTGLpSGQLQKAFVTGVP’ (SI=87.0%) is highly conserved in the LRR-VI-2 class (*N*m = 3523, *P* = 3523) and emerges in streptophyte algae (Fig. S5A). The LRR-VI-2 and LRR-Xb-1 motifs exhibit ubiquitous species presence across virtually all plant groups (from 490 plant species), suggesting the evolutionary conservation of this phosphorylation event. Nevertheless, the two lower-scored motifs observed in PERK-2 class appear in much fewer species (446 and 454 plant species, respectively) when compared with the motif observed in the LRR-VI-2 and LRR-Xb-1 classes. In addition, a few motifs with experimentally determined phosphorylation sites are found on several key RLK classes involved in plant development and defense of abiotic and biotic stimuli, including CrRLK1L-1, LRR-XI-1, RLCK-XII, and RLCK-VII (Ogawa et al., 2008; Tang et al. 2008; Chuang et al. 2011; Zhu et al., 2021). The motif ‘pTGpSpYApSpSLPpSBLCR’ from the CrRLK1L-1 class is also observed in a widely diverse set of 479 species (Table 1).

**Table 1.**
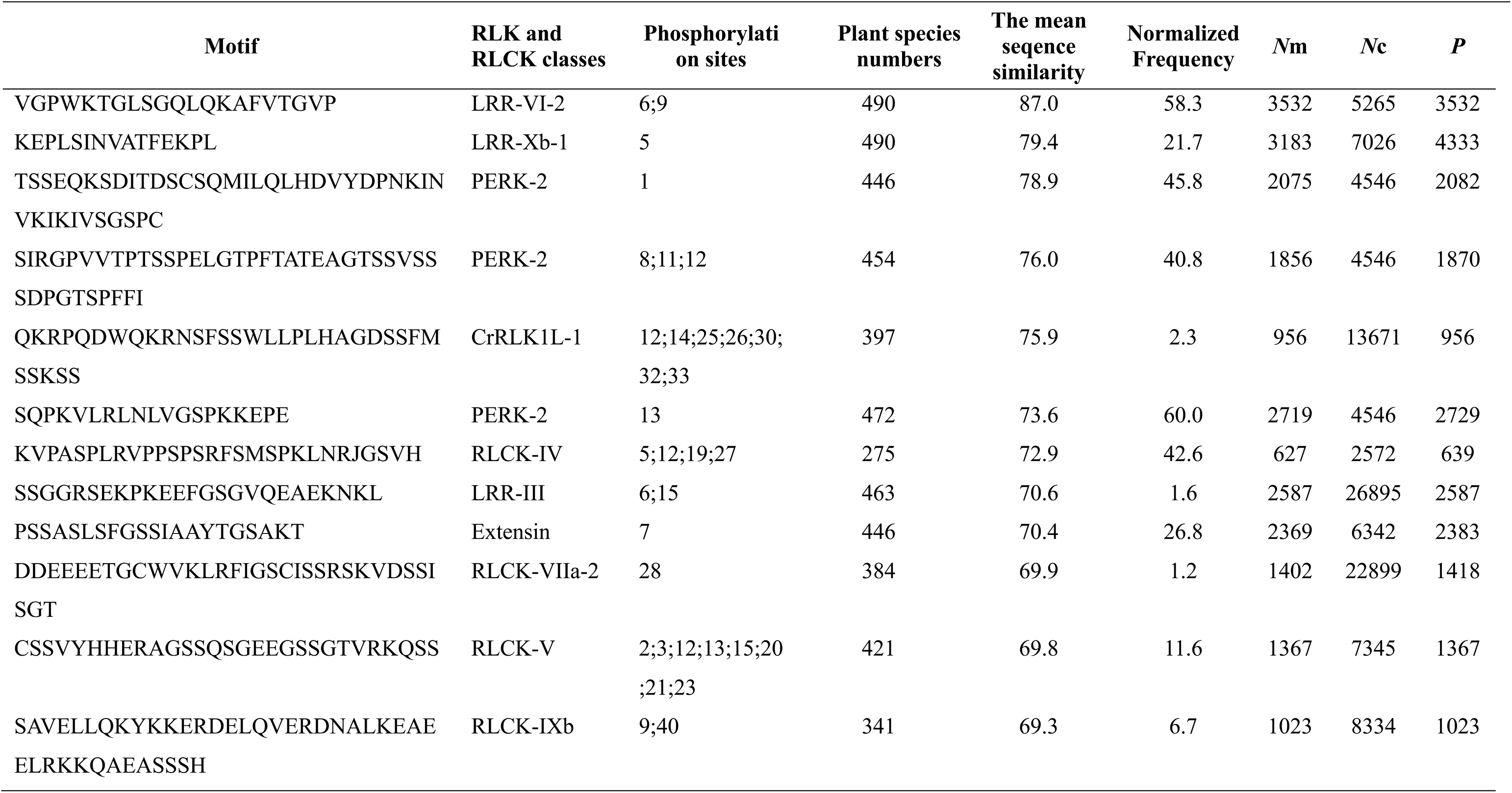
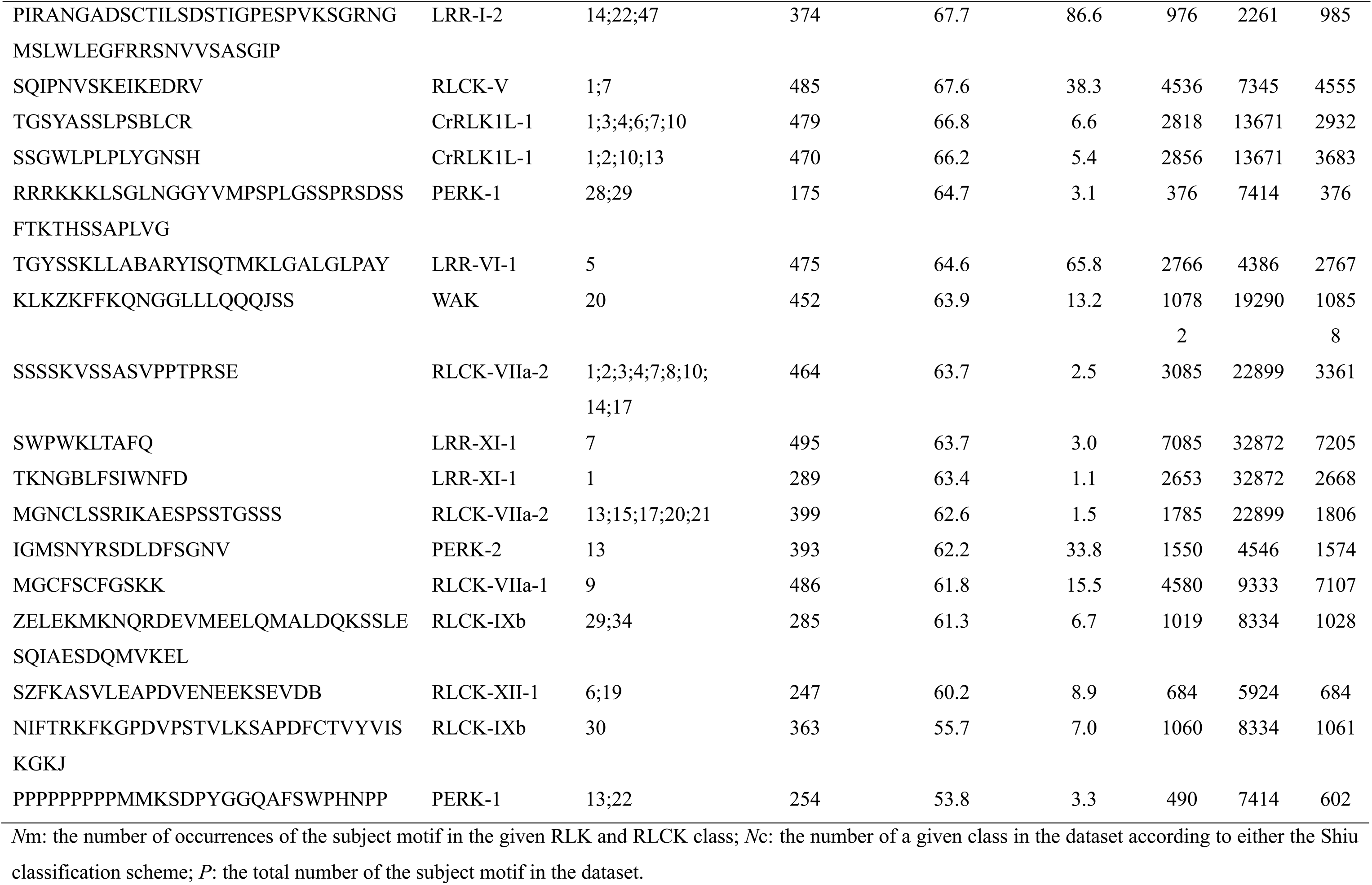
The identified 29 motifs in NRE segments with experimentally determined phosphorylation sites in classes from Shiu classification.

Out of the 65 analyzed motifs with a normalized frequency higher than 1, the other 32 motifs record no experimentally-determined phosphorylation event with similarity above 50%, while 28 still include at least one serine, threonine, or tyrosine that may potentially be phosphorylated (Table 2). These motifs also emerge in streptophyte algae (Fig. S5). The three highest-scored motif motifs ‘LSRNAPPGPPPLCSICQHKAPVFG’, ‘VVVAVKASKEIPKTALVWALTHVVQPGDCITLLVVVPSHSS’, and ‘GAVAAEAKRAQANWVVLDKQLKHEEKRCMEELQCNIVVMKR’ (SI ≧ 80%) are highly conserved in PERK-2 class.

**Table 2.** The identified 32 motifs in NRE segments without experimentally determined phosphorylation sites in classes from Shiu classification.

| Motifs | RLK and RLCK classes | Plant species numbers | The mean sequence similarity | Normalized Frequency | Nm | Nc | P |
| --- | --- | --- | --- | --- | --- | --- | --- |
| LSRNAPPGPPPLCSICQHKAPVFG | PERK-2 | 489 | 82.6 | 69.2 | 3134 | 4546 | 3144 |
| VVVAVKASKEIPKTALVWALTHVVQPGDCITLLVVVPSHSS | PERK-2 | 467 | 80.4 | 55.5 | 2513 | 4546 | 2522 |
| GAVAAEAKRAQANWVVLDKQLKHEEKRCMEELQCNIVVM<br>KR | PERK-2 | 480 | 80.0 | 61.4 | 2782 | 4546 | 2792 |
| PQEIFFDVP AEEDPEVHLGQL | LRR-II | 485 | 78.6 | 16.8 | 3658 | 9996 | 3658 |
| RSGAPQKVLPIEI | RLCK-VIII | 431 | 78.2 | 12.4 | 930 | 5635 | 1004 |
| LWGFPRFAGDCASGH | PERK-2 | 470 | 78.1 | 52.1 | 2364 | 4546 | 2374 |
| MRRWLCCTCQV | RLCK-VIII | 442 | 77.9 | 24.3 | 1694 | 5635 | 1705 |
| ITAPSPLVGLPEFSHLGWGHW | RLCK-V | 486 | 76.9 | 35.7 | 4218 | 7345 | 4224 |
| DVEAEMERLRLELKQTMDMYSTACKEALDAKQ | RLCK-IXb | 350 | 75.0 | 7.1 | 1080 | 8334 | 1088 |
| MFCCGGAE EEPYGPPANQYTAPP | RLCK-VIII | 387 | 72.4 | 11.1 | 800 | 5635 | 832 |
| DIHMVEAGNMV | LRR-IX | 466 | 70.6 | 51.0 | 1983 | 4182 | 2018 |
| GVLHRVTHPMGYQTKACPSF | PERK-2 | 246 | 70.3 | 9.2 | 422 | 4546 | 427 |
| DGRLSTDQAKEFYRKSASPLVSLE | LRR-IV | 383 | 69.3 | 127.0 | 1220 | 2096 | 1221 |
| NGSILLEKLIASCNGKSNPIR | RLCK-XII-2 | 164 | 69.0 | 150.7 | 1210 | 1678 | 1579 |
| GTSIRAMEEEVSKKVDAYVNMLLQSAEECEDEGV SIEVKI | PERK-2 | 216 | 68.6 | 8.0 | 365 | 4546 | 370 |
| YAEVREDWELEYGPH | L-LEC | 473 | 68.4 | 7.4 | 8543 | 23040 | 8560 |
| WNPKS RVMGSTRKEVT VFTDI | LRR-XV | 321 | 68.2 | 95.3 | 693 | 1823 | 694 |
| MGCRC SKLSPC | RLCK-XII-1 | 464 | 68.0 | 38.9 | 3064 | 5924 | 3149 |
| SSDTESENLSLSSASLRFQPWITEYLSS | PERK-2 | 293 | 67.5 | 24.3 | 1104 | 4546 | 1113 |
| LLWALQNFAGKKICILHVHQ | RLCK-IXb | 389 | 65.8 | 10.9 | 1656 | 8334 | 1661 |
| TIPKRDDHEIDMSRLDVDSMPPPPPPPP | LRR-V | 257 | 65.7 | 7.1 | 764 | 6996 | 764 |
| YDQLEQAMAEAEENSRRFAFEESVRRRKAE | RLCK-IXb | 421 | 64.8 | 9.1 | 1390 | 8334 | 1392 |
| PAQMIPMMGAKFPASSLKEZZVRAYREIE | RLCK-IXb | 348 | 63.7 | 5.8 | 882 | 8334 | 882 |
| YSRKMKDLKSKKAIYVCZQAPPSCHIWFIKGHLYTREA | RLCK-IXb | 449 | 63.6 | 14.0 | 2132 | 8334 | 2135 |
| PSRVVIAYDATKDRNEHELKLTIDNIRMGRGDILRGDTLLV | PERK-2 | 223 | 63.0 | 7.9 | 361 | 4546 | 365 |
| KEDTSSPVGVLEDYFRSSDSETSSSKEPTSDSESQQNSKPASR | RLCK-VI | 165 | 62.8 | 2.6 | 339 | 7655 | 339 |
| WHG |  |  |  |  |  |  |  |
| ENXCVQESHLLPVKDKSSEKDSG | RLCK-V | 51 | 60.3 | 1.7 | 198 | 7345 | 198 |
| YLLICARKKVQAEKVVIEMDDVAKGIVELISQHGITKLVMG | RLCK-IXb | 447 | 59.7 | 13.2 | 2010 | 8334 | 2012 |
| AA |  |  |  |  |  |  |  |
| QPTNQMEKVPKEAVVVKPDGHQTESRRMGAIPKPQBEQE | LRR-V | 99 | 58.4 | 2.7 | 293 | 6996 | 293 |
| LSTSQIIQKQNKSVFRQKSSEAPJLCAACGLRTELY | PERK-2 | 128 | 58.2 | 4.7 | 215 | 4546 | 217 |
| RHLEMYDKIEKFLQDYGNLKP | LRK10L-2 | 226 | 56.4 | 5.3 | 2476 | 14630 | 2476 |
| LEAJRKAKAAESLAEEEEAKQRKEAEEALAK | RLCK-IXb | 394 | 55.5 | 9.5 | 1450 | 8334 | 1460 |

Nm: the number of occurrences of the subject motif in the given RLK and RLCK class; Nc: the number of a given class in the dataset according to either the Shiu classification scheme; P: the total number of the subject motif in the dataset.

### Sequence features of NRE segments from LRR-VI-2 class

To explore the importance of the above phosphorylation motif in LRR-VI-2, we further analyze NRE sequence features, revealing that the NRE segment of a subset of LRR-VI-2 is astonishingly conserved in amino acid composition. This sequence comprises three regions: an N-terminal uncharacteristic conserved region, a phosphorylation site region, and a C-terminal α-helix characteristic to all RLKs (Fig. 5A). The phosphorylation site region features a conserved determined experimentally serine phosphorylation site located at −28 counted from the C terminal to the N terminal.

**Figure 5.**
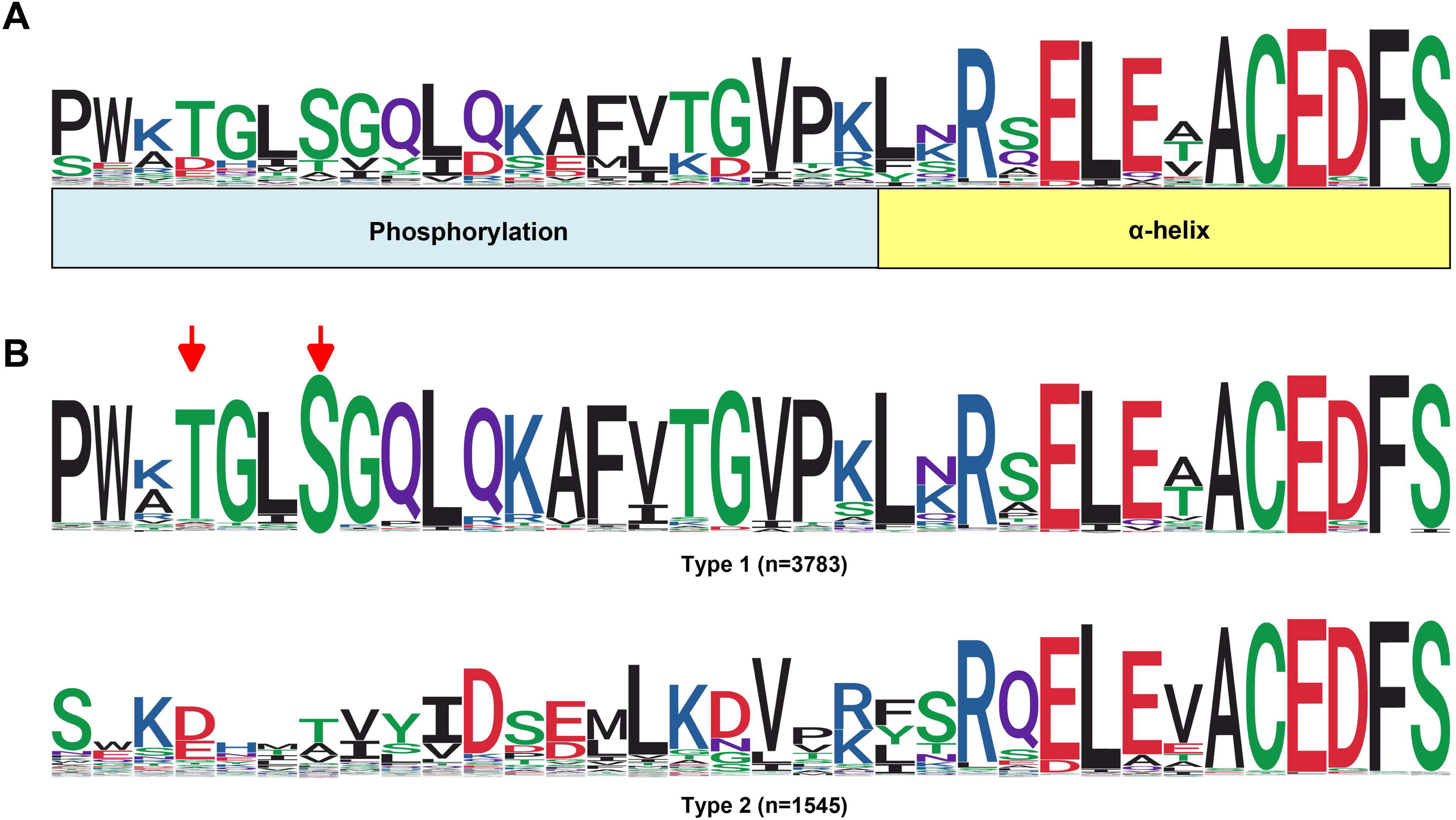
Sequence features of NRE segments in LRR-VI-2 class. (**A**) A schematic of NRE sequence characterized in LRR-VI-2 class. (**B**) Two types from the NRE segment in the LRR-VI-2 class; 1) Type 1 with two experimentally determined phosphorylation sites at −28S and −31T; 2) Type 2 refers to LRR-VI-2 with no apparent phosphorylation site at −28S and −31T. The red arrow indicates the phosphorylation sites.

WebLogo analysis of the LRR-VI-2 NRE sequence reveals two major types according to the presence or absence of the serine at −28 site; each exhibits distinctive features (Fig. 5B). LRR-VI-2 with a single phosphoserine site at −28S (n=3783) is the most abundant type. An experimental phosphothreonine site at −31T also often occurs in this type. However, the other type of NRE segments that exhibit no apparent serine phosphorylation site at position −28 or a putative threonine phosphorylation site are also observed (n=1545).

### The NRE phosphorylation of LRR-VI-2 protein MDIS1 affects its interaction with the co-receptor MIK1

Multiple sequence alignment of the kinase domain reveals that several characteristic kinase residues are not conserved in members of the LRR-VI-2 class (Fig. S6), rendering it a group of pseudokinases with compromised kinase activity. An LRR-VI-2 protein MDIS1 is known to form a heterodimer with MIK1 on the pollen tube cell surface that perceives female attractant LURE1 peptide ligand (Wang et al., 2016). Therefore, we select MDIS1-MIK1 interaction as a model system to verify the functional relevance of NRE phosphorylation in the LRR-VI-2 class.

The cytoplasmic regions of MDIS1 (MDIS1^CD^) and MIK1 (MIK1^CD^) are recombinantly constructed using His tag and GST tag for affinity chromatography, respectively. Both recombinant proteins are expressed in bacteria and purified (Fig. S7). Interactions between MDIS1^CD^ and MIK1^CD^ are evaluated using GST pull-down assays. The His-MDIS1^CD^ WT interacts with the GST-MIK1^CD^, but not with the GST protein (Fig. 6A). While the His-MDIS1^CD^ S350A mutant exhibits comparable binding affinity towards GST-MIK1^CD^, His-MDIS1^CD^ S350D mutant exhibits much stronger affinity than either His-MDIS1^CD^ WT or His-MDIS1^CD^ S350A towards GST-MIK1^CD^.

**Figure 6.**
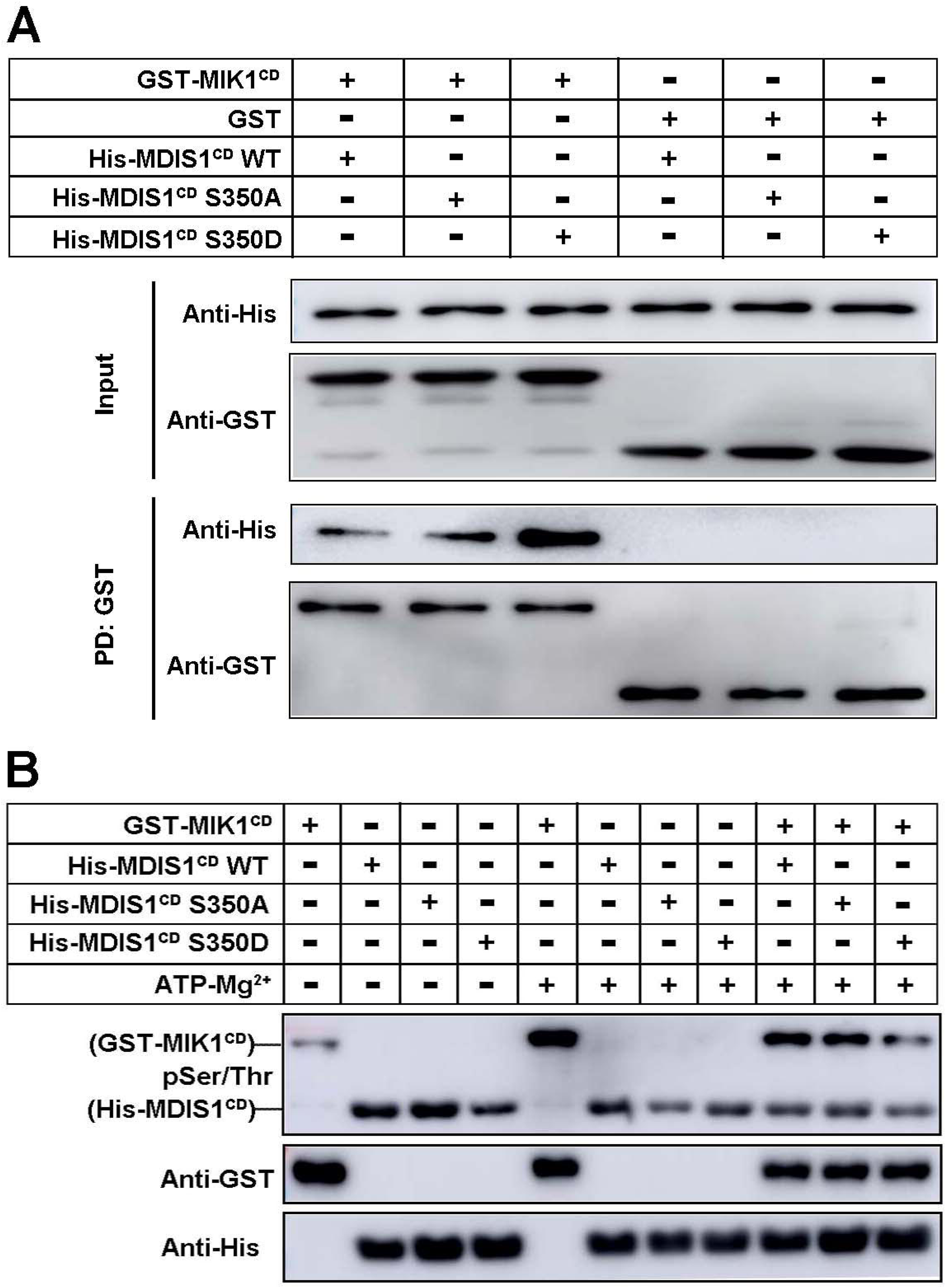
The phosphorylation site of NRE segments of *Arabidopsis* LRR-VI-2 MDIS1 plays a key role in MIK1 kinase activity. (**A**) GST pull-down assays. GST-MIK1^CD^ pulled down the tested His-MDIS1s. Proteins in the upper panel are detected using an anti-His antibody, and those in the lower panel are detected using an anti-GST antibody. (**B**) *in vitro* kinase assays. Proteins in the upper panel are detected using an anti-pSer/Thr antibody, and those in the lower panel are detected using anti-GST and anti-His antibodies.

### MDIS1 NRE phosphorylation alleviates the auto-phosphorylation of its co-receptor MIK1

We further investigate the functional role of the phosphorylation site using *in vitro* kinase assays. Strong auto-phosphorylation of GST-MIK1^CD^ is detected, whereas His-MDIS1^CD^ self-phosphorylation activity is weaker (Fig. 6B). Incubation of GST-MIK1^CD^ together with His-MDIS1^CD^ S350D mutant results in marked reduced phosphorylated level for GST-MIK1^CD^ protein (Fig. 6B). By contrast, GST-MIK1^CD^ proteins exhibit a similar phosphorylation pattern when incubated with either His-MDIS1^CD^ WT or His-MDIS1^CD^ S350A mutant.

## Discussion

### A 14-amino acid motif forms a characteristic α-helix upstream of the core RLK and RLCK kinase domain

A consensus motif ‘FSYEELEKATBNFS’ located at the C-terminal of NRE segments appears to be characteristic in the majority of RLK and RLCK sequences (Fig. 3D-F), with the exceptions in only nine classes of RLK superfamily (LRR-III, LRR-VI-2, LRR-VII-1, LRR-XI-2, LRR-XIIIa, LRR-XIIIb, RLCK-II, Singleton, and URK-2) out of 65 RLK and RLCK classes defined in the Shiu classification scheme. The C-terminal of NRE segments in these nine classes of the RLK superfamily, albeit deviate from the consensus motif, exhibit highly characteristic features within the same classes yet appear to be mutually distinctive among different classes (Fig. 7A), indicating the possible presence of a diversified profile of alternative regulatory mechanisms that deserve further exploration.

**Figure 7.**
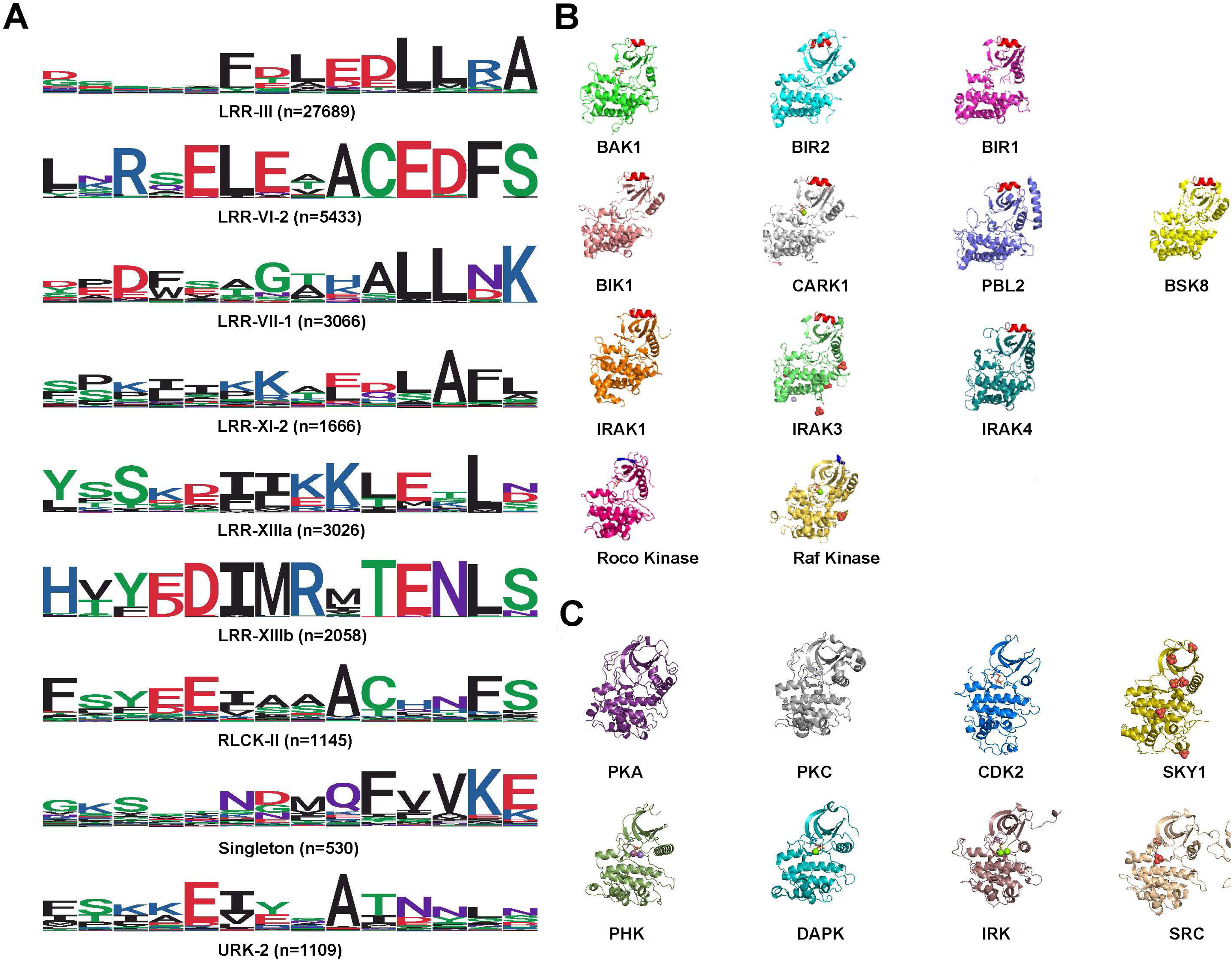
The sequence features and structure of the C-terminal of NRE segments in RLKs. (**A**) Sequence features of C-terminal of NRE segments from nine classes of RLK superfamily in which position −5 is not T or position −2 is not F. (**B**) The structures of representative RLK, RLCK, and protein kinases; row 1: RLKs including BAK1 (3TL8), BIR2 (4L68), BRI1 (4OH4); row 2: RLCKs including BIK1 (5TOS), CARK1 (5xd6), PBL2 (6J6I), BSK8 (4I92). row 3: IRAKs including IRAK1 (6BFN), IRAK3 (6RUU), IRAK4 (6TI8); row 4: Roco kinase (4F0F), Raf kinase (6U2G). (**C**) The structures of non-RLK protein kinases with the universal absence of the RLK-specific α-helix. row 1: AGC kinases (PKA: 2CPK; PKC: 2JED), CMGC kinases (CDK2: 1FIN; SKY1: 1HOW); row 2: Calcium/calmodulin kinases (PHK: 2PHK; DAPK: 1JKK), Tyrosine kinases (IRK: 1IR3; SRC: 1Y57). Molecular graphics are prepared using PyMOL (DeLano Scientific). Molecular surfaces are rendered with a probe radius of 1.4 Å.

This characteristic motif ‘FSYEELEKATBNFS’ forms an α-helix located right above the upper lip of the core kinase domain (Fig. 3J, K). The presence of this α-helix has been reported as an extra N-terminal portion (G272-N291) that forms a long extension and caps the top of the twisted β-sheets in the N-lobe of the core kinase domain in the RLK BAK1 (Yan et al., 2012). The universal presence of the α-helix is further illustrated by its presence in a series of RLK structures (Fig. 7B), including the AvrPtoB’s host RLK BAK1 (3TL8) (Cheng et al., 2011), BIR2 (4L68) (Blaum et al., 2014), and the BKI1 inhibiting protein BRI1 (4OH4) (Wang et al., 2014). In addition, some RLCKs having NKE segments structures also reveal the same sequence motif and the same α-helix (Fig. 7B). For example, the structure of plant immunity-related BIK1 (5TOS) (Lal et al., 2018) is an RLCK that lacks the TM region, yet still possess this characteristic upstream α-helix. Structural comparison of plant RLKs with the human IRAKs (Interleukin 1 (IL-1) receptor-associated kinases) reveals that IRAK1 (6BFN) (Wang et al., 2017), IRAK3 (6RUU) (Lange et al., 2021), and IRAK4 (6TI8) (Degorce et al., 2020) all possess a similar α-helix located right upstream of the core kinase domain (Fig. 7B). Multiple sequence alignments indicate characteristic features at the sequence immediately upstream of the N-terminal in IRAK1, IRAK3, and IRAK4, as we observed ‘GTHNFS,’ ‘GTRNFH,’ and ‘VTNNFD’ motifs, respectively.

Non-RLK plant kinases also contained a core kinase domain. We find a ‘KQKVGELKDDDFE’ motif in MKK1 (MAPK Kinase) ERK activator and a ‘KSRLPTLADNEIE’ motif in Ras of complex protein. These two sequences appear distantly related to the characteristics of the plant RLK superfamily motif ‘FSYEELEKATBNFS.’ However, close examination of either the Roco kinase (4F0F) (Gilsbach et al., 2012) or the Raf kinase (6U2G) (Liau et al., 2020) structure fails to identify the presence of α-helix located upstream of the core kinase domain (Fig. 7B). Close examination of structural representatives from other protein kinase families indicates the universal absence of the RLK-specific α-helix in any other plant kinases (Fig. 7C) (Knighton et al., 1991; Jeffrey et al., 1995; Hubbard, 1997; Lowe et al., 1997; Nolen et al., 2001; Tereshko et al., 2001; Cowan-Jacob et al., 2005; Jacobs et al., 2006).

To explore the functional significance of the α-helix motif of the RLK superfamily, we focus on the FER protein, which is heavily involved in plants development and stress response (Stegmann et al., 2017; Hansen et al., 2019; Chen et al., 2020; Zhang et al., 2020a), and participates in a wide array of physiological processes (Zhu et al., 2021). FER possesses a typical cytoplasmic kinase domain (KD) and exhibits both kinase and auto-phosphorylation activities (Li et al., 2018), yet the regulation of its kinase activity by NRE remains inconclusive. Our data suggest that the presence of the α-helix of the NRE segment leads to the repression of FER kinase activity (Fig. 4). Besides its role in regulating the kinase activity, the C-termini containing α-helix of the NRE segments are also found essential for chitin response in both *At*CERK1 and *Os*CERK1 (Zhou et al., 2020). To sum up, the characteristic α-helix forming an N-terminal extension of the core kinase domain is a hallmark of the RLK superfamily, while “FSYEELEKATBNFS” and some of its variations with minor classes define the N-terminal boundary of the core kinase domain.

### Potential phosphorylation sites in the NRE segments

Phosphorylation is a reversible modification catalyzed by RLK, which transfers a phosphoryl group from ATP, particularly to the hydroxyl group of specific serine, threonine, or tyrosine residues within their target proteins (Champion et al., 2004). However, most of these studies mainly focus on phosphorylation in the kinase region, and few have looked at the NRE segment. Herein, we report the most enriched and comprehensive phosphorylation motif data in the NRE segment. These motifs are further associated with RLK and RLCK classes to aid in targeted phosphorylation analysis of specific sets of proteins and provide a strong foundation and reference to infer the conservation of phosphorylation sites in NRE segment across plant species.

Generally, the NRE segment possesses a high serine content compared with other parts of the RLK and RLCK sequence. The rich presence of serine residues indicates that NRE segments are at the forefront of the region where frequent phosphorylation events occur (Fig. 2 D-F). Throughout the phosphorylation motif analysis using experimental data cataloged by the EPSD databases, some motifs are specific in a few particular RLK or RLCK classes, while other motifs distribute widely in a diverse set of RLK or RLCK classes (Table S3, Table 1). Phosphorylation mainly occurs on serine and threonine residues in plants. However, the relative frequency of tyrosine phosphorylation is around 5%, which is no less than that in humans, implying that plant signal transduction could also rely on tyrosine phosphorylation despite the lack of animal receptor tyrosine kinase (RTK) (Nakagami et al., 2010; van Wijk et al., 2014). An example is the brassinosteroid (BR) receptor kinase BRI1, whose juxtamembrane segment is confirmed as an activator kinase domain and determinant of autophosphorylation specificity (serine/threonine relative to tyrosine) (Oh et al., 2012). Moreover, a few key RLK classes are enriched in conserved motifs with experimentally determined phosphorylation sites. The plant-specific CrRLK1L-1 is an important RLK class in plant development and defense, which participate in widespread cellular processes and is receiving increasing attention as a regulator of cell expansion, plant reproductive development, and immunity (Zhu et al., 2021). In our study, we found that the NRE segment of the CrRLK1L-1 class is enriched with the motif ‘pTGpSpYApSpSLPpSBLCR,’ which similarity is above 66.8% and widely occurs in 479 species. Especially in addition to serine/threonine phosphorylation, this motif also has tyrosine phosphorylation, demonstrating that the CrRLK1L-1 class may be a dual-specificity kinase class. The highly conserved phosphorylation motif in NRE segement that features multiple serines and phosphotyrosine is likely to play a significant regulatory role in the transduction of defense signals from cell-surface receptors to downstream signaling components in plants for the CrRLK1L-1 class.

The motif ‘VGPWKpTGLpSGQLQKAFVTGVP’ that exists in the specific LRR-VI-2 class coincides with the reported phosphorylation site and is the most conserved and universal to plant species. The presence of highly conserved phosphorylation sites across multiple species indicates the ubiquitous roles of these sites in regulating the plant signal transduction pathway. Further investigation of the consensus sequences indicates highly conserved features around two phosphorylation sites (−28S and −31T) in LRR-VI-2 class (Fig. 5), inferring a possible synergistic mechanism of two phosphorylation sites. Our motif analysis also identifies some novel and conserved motifs having S/T/Y residues specific to the RLK and RLCK classes with no reported NRE phosphorylation events. In short, characterizing NRE segment sequence motifs with phosphorylation sites can help build a predictive algorithm that can distinguish S/T/Y residues prone to phosphorylation from those disincline to phosphorylation in the NRE segments in RLKs and RLCKs.

### The schematic structure of a ‘kinase switch’ in the LRR-VI-2

Our schematic functional division of the NRE segment in LRR-VI-2 provides a paradigm that applies to most NRE segments (Fig. 5A). While most NRE segments are considered intrinsically disordered and are liable to significant sequence variability, the NRE segment of one group of LRR-VI-2 is astonishingly conserved in both length and composition. The length of this conserved sequence is also at the peak length of a typical NRE sequence (Fig. 2A).

One can speculate that the single phosphorylation site located upstream of the C-terminal α-helix characteristic to all RLKs could behave like an on/off switch to alternate the conformation of both the C-terminal α-helix’s and hence the downstream core (pseudo)-kinase domain (Hornbeck et al., 2015). For example, an unphosphorylated NRE segment of RLK ACR4 in *Arabidopsis* may preferentially interact with a kinase domain near the activation loop, presumably holding the kinase domain in an inactive state. Phosphorylation of this NRE segment was found to preferentially interact with the N-terminal lobe of the kinase domain, resulting in an ACR4 variant with a defect in substrate phosphorylation (Meyer et al., 2013). RLK BAK1 was reported to transform CERK1 from an off state into a primed state by inducing allosteric change by mediating NRE phosphorylation (Gong et al., 2019). Interestingly, the motif “VGPWKTGLSGQLQKAFVTGVP” characterized from LRR-VI-2 based on a total of 3532 NRE segments from 490 plant species is thus far the most conserved and contains an experimentally-determined phosphorylation site “TGL(pS)GQLQK” referring to −28 site (Table 1, Fig. 5B).

Members of the LRR-VI-2 are atypical kinases (Castells and Casacuberta, 2007), with variations in the glycine-rich loop and the VAIK/HRD/DFG motifs in the catalytic domain that are essential for catalysis (Fig. S6). A well-studied member of the LRR-VI-2 class is MDIS1, which forms a receptor heteromer with MIK1 on pollen tube cell-surface that perceives female attractant LURE1 (Wang et al., 2016). Binding assay suggests the stronger binding between MDIS1 and its co-receptor MIK1 using aspartic acid mutant S350D as a phosphorylation mimicry (Fig. 6A). The presence of the exact phosphorylation mimicry also leads to the compromise of kinase activity (Fig. 6B), likely through allosteric regulation or scaffolding molecules in cell signaling (Zeqiraj and van Aalten, 2010; Reiterer et al., 2014; Kung and Jura, 2019). Allosteric regulation might require dynamic transitions between different conformational states as seen for canonical kinases (Huse and Kuriyan, 2002; Kornev and Taylor, 2015). Alternatively, pseudokinase domains might be more static than other kinase domains, allowing them to serve as rigid scaffolds. The common presence of serine in the LRR-VI-2 pseudokinases subclass suggests its crucial role in the activation process despite the variations of essential residues in the kinase domain. These results indicate that the phosphorylation site (position −28) of the NRE segment of *Arabidopsis* LRR-VI-2 MDIS1 is necessary to regulate MIK1’s kinase activity.

The relatively short NRE segments provide a minimal set of functional elements that coordinate with each other to complete its task as a “kinase switch.” While the characteristic C-terminal α-helix is universal to all RLKs and RLCKs, this LRR-VI-2 model suggests a plausible mechanism through which phosphorylation regulates downstream kinase activities in RLK. The “kinase switch” may control the kinase activity either directly or allosterically through its binding partners upon phosphorylation of the NRE segment.

## Materials and Methods

### Plant genomes used for RLK and RLCK sequences identification

Plant genomes containing RLK and RLCK for NRE analysis are obtained from several sources. The sources include genomes of 84 plant species from the Joint Genome Portal (JGI, https://genome.jgi.doe.gov/portal/) (Grigoriev et al., 2012); 55 species from The Genome Warehouse (GWH) (Chen et al., 2021); 45 species from Ensembl Plants (Yates et al., 2022); 53 species from the NCBI database (https://www.ncbi.nlm.nih.gov/); 29 species from the Amazon Web Services (AWS) (https://uc3-s3mrt1001-prd.s3.us-west-2.amazonaws.com/); 32 species from the GigaDB database (http://gigadb.org/site/index); 30 species from the ePlant database (http://eplantftp.njau.edu.cn/); 14 species from the Medicinal Plant Genomics Resource (MPGR, http://mpgr.uga.edu/); 14 species from the Hardwood Genomics Project (HGP, https://www.hardwoodgenomics.org), and another 172 species from other databases. References and relevant information regarding this non-redundant set of 528 species and their source are available in Supplemental Data (Table S1). The gene domain organization and gene structure for plant genomes from GenBank and RefSeq in NCBI is predicted using the EST-based method or transcript-to-genome sequences. In addition, some recently-published genomes also use *de novo* prediction and homology-based methods for prediction. The 528 species are divided into seven types (Algae, Charophytes, Bryophytes, Lycophytes, Ferns, Gymnosperms, and Angiosperms). Each includes one or more clades, with Algae containing red algae, brown algae, and green algae; Bryophytes containing liverworts and mosses; Gymnosperms containing ginkgo and conifers; and Angiosperms containing basal angiosperms, monocots, and eudicots.

### Identification of NRE sequences in RLK superfamily

RLKs and RLCKs sequences are identified and classified from the genome sequences of the 528 plant species using a set of pre-compiled HMMs (Lehti-Shiu and Shiu, 2012) as implemented in the iTAK program (Zheng et al., 2016). RLKs and RLCKs designation and classification are obtained using the latest Shiu classification scheme according to the plant protein kinase phylogeny and extracellular domain identities (Lehti-Shiu and Shiu, 2012). In addition, the transmembrane region for each RLK and RLCK sequence is identified as a consensus of the outputs of the TMHMM (Sonnhammer et al., 1998; Krogh et al., 2001) and Phobius (Käll et al., 2004) analysis. The N-terminal boundary of each NRE segment is defined as the residue that immediately follows the C-terminal residue of the transmembrane sequence. The boundary of the cytoplasmic kinase domain is identified using Pfam (El-Gebali et al., 2019), with two Pfam families, PF00069 referring to “PKinase” and PF07714 referring to “PK_Tyr_Ser-Thr.” Inspection of the HMM profiles for both Pfam families indicates a characteristic (L/I/V)GXG motif between positions 5-8 at the N-terminal of the kinase domain. We define the NRE C-terminal boundary immediately before the N-terminal residue of the kinase domain. All NRE sequences and the associated information are imported into a custom PostgreSQL database for further analysis.

### Analysis of NRE sequences

The length and composition of the NRE are calculated using a custom python script which allows plotting either the whole dataset or its subset according to the RLK/RLCK classes or the types of plant species. WebLogo (Schneider and Stephens, 1990) analyzes the specific distribution of the amino acid profiles at each position. Amino acid profiles are constructed using the first 20 amino acids of the NRE aligned to the N-terminal, and the last 20 amino acids of the NRE aligned to the C-terminal. Hydrophobic, polar, acidic, basic, and neutral residues are shown in black, green, red, blue, and purple in the WebLogo profile, respectively.

### ATPase activity assay

The cytoplasmic domain of FERONIA (FER) (residues 518-816) wild type and its corresponding mutant (residues 518-816, K565R) are cloned into modified pRSFDuet-1. The kinase domain of FER (residues 536–808) wild type not containing 14-amino acids in the C-terminal of the JM segment is cloned into pET-28a. Recombinant proteins are expressed and purified using a protocol described previously (Chen et al., 2016). Kinase-Lumi^TM^ Luminescent Kinase Assay Kit (Beyotime, China) is used in ATPase activity assay. Specifically, 1 μM proteins are incubated in 50 μL kinase reaction mixture containing 50 mM HEPES buffer, pH 7.5, 10 mM MgCl_2_, 10 mM MnCl_2_, 1 mM EGTA, 10 μM ATP, followed by the addition of 50 μL Kinase-Lumi Chemiluminescence Kinase Detection Reagent. Following reaction at room temperature for 10 minutes, chemiluminescence is detected using a multiscan spectrum (Thermo scientific, Fluoroskan Ascent^TM^ FL). The enzyme activity of variants and mutants of the FER cytoplasmic domain is deduced according to the standard curve using the measured FER protein substrate.

### Motif discovery and analysis

Based on the WebLogo analysis, 14 residues from the C-terminal are removed before motif analysis. Motif analysis is conducted with the middle part of the NRE segments using the MEME Suite, including MEME for motif discovery and FIMO for motif identification. MEME is used to predict the potential motif with a *p*-value threshold of 0.005 (Bailey and Elkan, 1994), while FIMO is used to identify the species sequences containing these motifs with a *p*-value threshold of 0.001 (Grant et al., 2011).

The normalized frequency of each motif is calculated using the formula 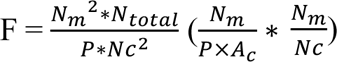, in which *N_m_* is the number of occurrences of the subject motif in the given class according to the Shiu classification scheme, *P* is the total number of the subject motif in the dataset, and *A_c_* is the abundance score of a given class in the dataset according to the Shiu classification scheme. The abundance score is calculated as 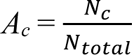, in which *N_c_* is the number of a given class in the dataset according to either the Shiu classification scheme or the types of plant species defined above, *N_total_* is the total number of NRE sequences in the dataset. Motifs with a normalized frequency higher than 1 are selected for further analysis. A normalized frequency higher than 1 indicates an overrepresented class for the given motif, while a normalized frequency lower than 1 indicates an underrepresented class for the given motif.

### Identification of phosphorylation sites in motifs of the NRE segments

Our current study uses experimentally determined phosphorylation data from 23 plant species to investigate the relationship between the NRE segment and phosphorylation motifs (Table S2). The 23 plant species genomes contain 35,913 NRE segments from 36,912 RLKs and RLCKs. Motifs identified using FIMO in the previous step are mapped onto the NRE segments in the RLK and RLCK sequences. A list of experimentally determined phosphorylation sites is obtained from the EPSD database (Lin et al., 2021). The starting and ending positions of the phosphorylation sites are compared within the boundary of FIMO motifs in the NRE segments to obtain a list of phosphorylation sites entirely contained in an identified motif. Motifs without serine/threonine/tyrosine residues at the consensus sequence at the expected phosphorylation site are excluded from further analysis. The consensus analysis of the identified motif sequence and the matching NRE sequence is investigated using the formula 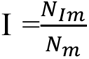, in which *N_Im_* is the number of consensus amino acids in the motif sequence and the NRE sequence, *N_m_* is the number of amino acids of the matching NRE sequence. The motif having more than 80% identity with the matching NRE sequences is considered highly conserved and similar. Motifs with sequence identity between 50% and 80% are considered moderately conserved and similar. The distributions of these motifs in various species are analyzed using R. In this analysis, 457,685 NRE segments without domain are analyzed except 67,263 NRE segments with domain.

### Sequence alignment

The representative sequences of LRR-VI-2 class cytoplasmic domains are retrieved from the abovementioned dataset. Multiple sequence alignment (MSA) is performed using MAFFT v7 (Katoh and Standley, 2013) with the L-INS-I settings, with figures prepared using Jalview 2.11.1.4.

### Expression and purification of MDIS1 and MIK1 protein

The cytoplasmic domains of MALE DISCOVERER1 (MDIS1, residues 335-641) and MDIS1-INTERACTING RECEPTOR LIKE KINASE1 (MIK1, residues 698-983) are cloned into modified pRSFDuet-1 and pGEX-6P-1 vectors containing an N-terminal 6×His tag and N-terminal glutathione S-transferase (GST) tag, respectively. We construct two MDIS1 mutants (S350A and S350D) to investigate the functional role of the phosphorylation site in the positions of serine −28 of the NRE segment. Mutants with aspartic acid (D) and alanine (A) at the −28 position are designed to mimic serine’s phosphorylation and dephosphorylation states, respectively. The fused proteins are expressed in the chemically competent *E. coli* BL21 (DE3) from TransGen Biotech. Cells are grown to an OD_600_ value of 0.6-0.8 at 37 °C and then induced with 0.5 mM Isopropyl β-D-thiogalactopyranoside (IPTG) followed by 18-20 h incubation at 16 °C. MIK1 is co-expressed with the λ-phosphatase for dephosphorylation. The cells are harvested by centrifugation at 4,200 rpm for 15 min, resuspended with 30 ml lysis buffer containing 20 mM Tris–HCl (pH 7.5), 150 mM NaCl, and 10 mM imidazole, and homogenized using a low-temperature ultra-high pressure cell disrupter (JNBIO, Guangzhou, China). The supernatants are collected after centrifugation at 12,000 rpm for 1 h at 4 °C. The His-MDIS1^CD^ protein is loaded onto a Ni-NTA agarose column (GE Healthcare) and eluted using elution buffer containing 20 mM Tris-HCl, 150 mM NaCl, pH 7.5, and 300 mM imidazole, and then exchanged into Tris buffer without imidazole. The GST-MIK1^CD^ is loaded onto GST agarose (Glutathione Sepharose 4B, GE Healthcare) and eluted using GST elution buffer containing 50 mM Tris–HCl, 20 mM glutathione, 150 mM NaCl, PH7.5.

### GST pull-down assay

Each of the purified His-MDIS1^CD^ wild-type (WT), His-MDIS1^CD^ S350D, His-MDIS1^CD^ S350A is mixed with purified MIK1-GST proteins and incubated with glutathione agarose beads overnight at 4°C in GST binding buffer (20 mM Tris-HCl, pH 7.5, 150 mM NaCl, 5% (vol/vol) glycerol). The beads are collected by centrifugation and washed five times with buffer containing 20 mM Tris-HCl, pH 7.5, 150 mM NaCl, 0.3% Triton X-100, 0.1% SDS. Finally, the proteins bound on the beads are boiled with 1× SDS/PAGE loading buffer in 95°C water bath and analyzed by both SDS-PAGE and immunoblotting with anti-GST (Abclonal, AE001) and anti-His (Abclonal, AE003) antibody.

### *in vitro* MDIS1 and MIK1 phosphorylation assays

For phosphorylated assays, each of the His-MDIS^CD^ WT, His-MDIS^CD^ S350D, and His-MDIS^CD^ S350A is co-incubated with GST-MIK1^CD^ at 25℃ for 1 h in assay buffer containing 25 mM Tris-HCl at pH 7.5, 10 mM MgCl_2_, 1 mM ATP. The reaction is stopped by adding 1× SDS loading buffer. The proteins are separated by 12% SDS-PAGE and immunoblotting. The phosphorylated GST-MIK1^CD^ and His-MDIS^CD^ proteins are detected using an antibody against pSer/pThr (ab9332, Abcam). In addition, the presence of the two recombinant proteins is verified by immunoblotting using either GST antibody or His antibody.

## ACKNOWLEDGMENTS

The authors would like to thank Shin-Han Shiu from Michigan State University and Jia Li from Lanzhou University College of Biology for the valuable discussions and comments. The authors would like to thank Marek Grabowski from the University of Virginia and Mingcong Chen for their help proofreading the manuscript. The authors thank Haojie Ma for preparing the RLK dataset for further analysis. This work was supported by grant 2021JJ30101 for Natural Science Foundation of Hunan, grant 2021RC2061 for the science and technology innovation Program of Hunan Province, grants 8197061423 and 3190080220 from the National Natural Science Foundation of China, startup funds provided by Hunan University, and a database construction fund from Hunan Haikun Co. Ltd.

## AUTHOR CONTRIBUTIONS

HZ designed the study; QF prepared the manuscript, conducted the molecular experiments and analyzed data; QL prepared the dataset and analyzed data; RZ performed the motif discovery analysis; JC conducted preliminary experiments on the FER kinase; JC, HG, and ZM provided essential discussion regarding the kinase activation mechanism presented in the study; HZ and FY supervised the study.

## AVAILABILITY OF MATERIALS

The 510,233 NRE sequences of RLK and RLCKs from 528 plant species and program codes supporting the findings of this study are available from GitHub (https://github.com).

## SUPPORTING INFORMATION

**Figure S1. Length distribution of NRE segments of different classes of plant RLK superfamily according to Shiu classification scheme and designation (65 classes).** (**A**) RLK sequences. (**B**) RLCK with TM region sequences. (**C**) RLK without TM region sequences.

**Figure S2. Composition distribution of EDQN residues of different classes of plant RLK superfamily according to Shiu classification scheme and designation (65 classes).** Red line refers to Aspartic acid (D), dark red line refers to Glutamic acid (E), green line refers to Asparagine (N), light green line refers to Glutamine (Q).

**Figure S3. Composition distribution of STY residues of different classes of plant RLK superfamily according to Shiu classification scheme and designation (65 classes).** Green refers to Serine (S), olive refers to Threonine (T), lime refers to Tyrosine (Y).

**Figure S4. Composition distribution of ALF residues of different classes of plant RLK superfamily according to Shiu classification scheme and designation (65 classes).** Dark line refers to Alanine (A), grey refers to Leucine (L), beige refers to Phenylalanine (F).

**Figure S5. The distribution of 61 motifs in 7 types of plants groups (Mean Similarity > 50%).** (**A**) The distribution of 29 motifs with experimentally determined phosphorylation sites. (**B**) The distribution of 32 motifs without experimentally determined phosphorylation sites.

**Figure S6. Multiple sequence alignment of the kinase domain in the four representatives of the LRR-VI-2 class and the MIK1 and BAK1 RLKs.** Roman numbers mark different subunits of the kinase domain. Asterisks on the top designate the residues conserved in active kinases. The four boxes I, II, VIb, and VII highlight critical catalytic residues (glycin-rich loop, VAIK motif, HRD and DFG motif) that LRR-VI-2 class lacks.

**Figure S7. The images of MIK1 and MDIS1 fusion protein gels.** Four fusion proteins are expressed and purified before loaded for SDS-PAGE and stained with Coomassie brilliant blue G-250.

**Table S1.** Source of the 528 plant species considered in the current study

**Table S2.** The information of 23 plant species with experimentally determined phosphorylation sites

**Table S3.** 31 motifs with experimentally determined phosphorylation sites in NRE segments

