## Supplemental file for "Large-scale analysis of the N-terminal regulatory elements of the kinase domain in plant receptor-like kinase family"

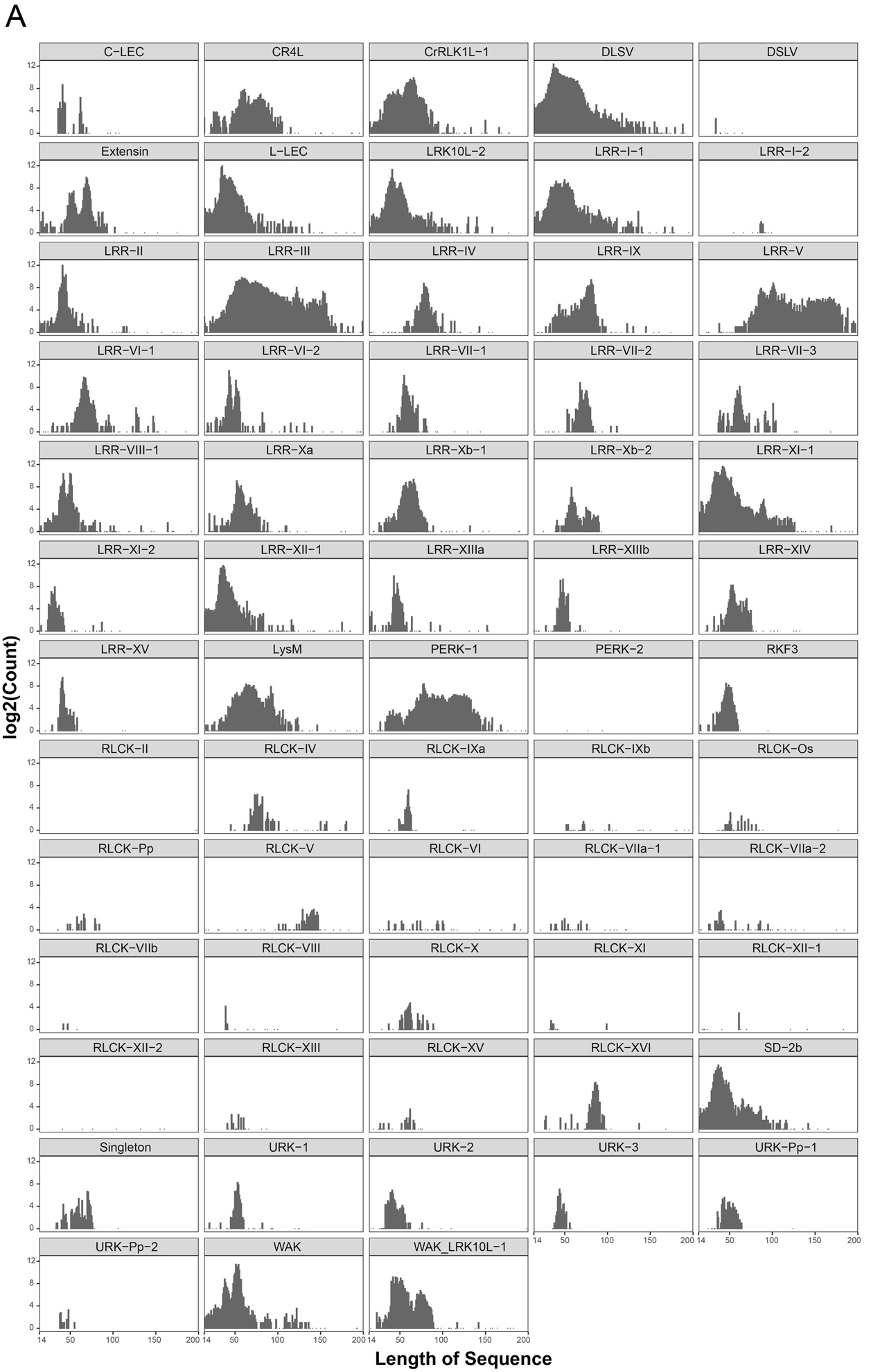

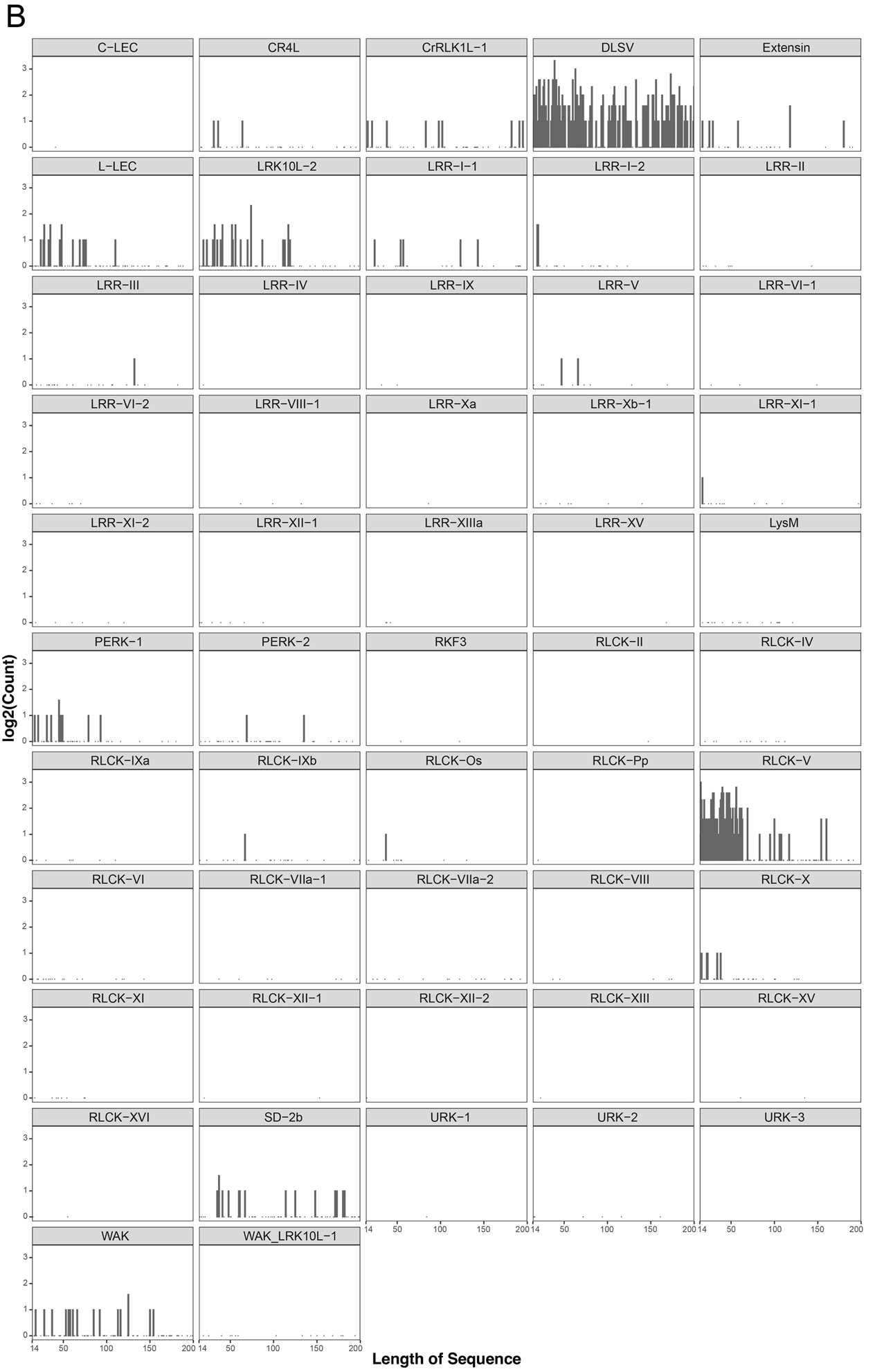

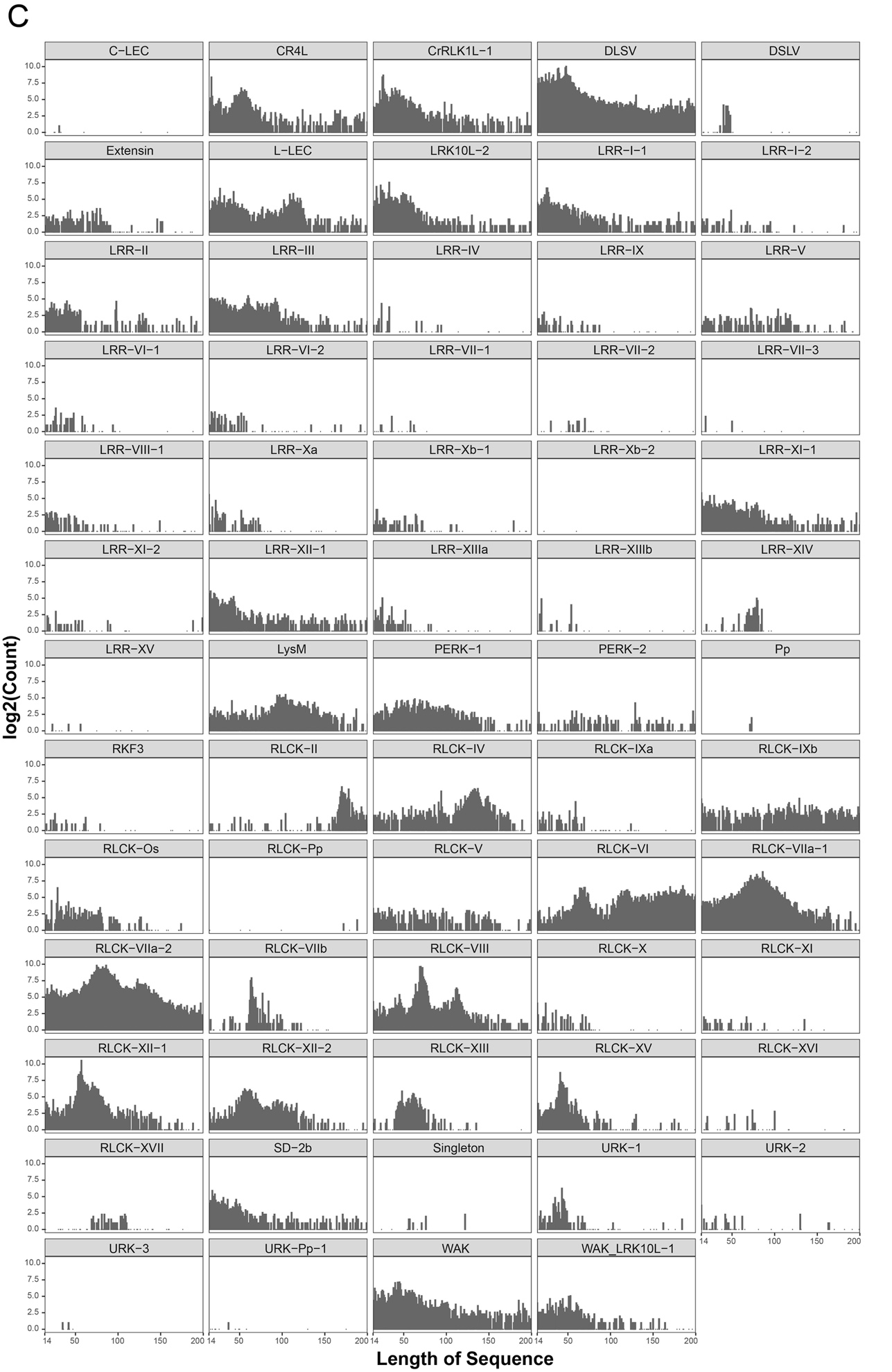

**Figure S1. Length distribution of NRE segments of different classes of plant RLK superfamily according to Shiu classification scheme and designation (65 classes).**

(**A**) RLK sequences.

(**B**) RLCK with TM region sequences.

(**C**) RLK without TM region sequences.

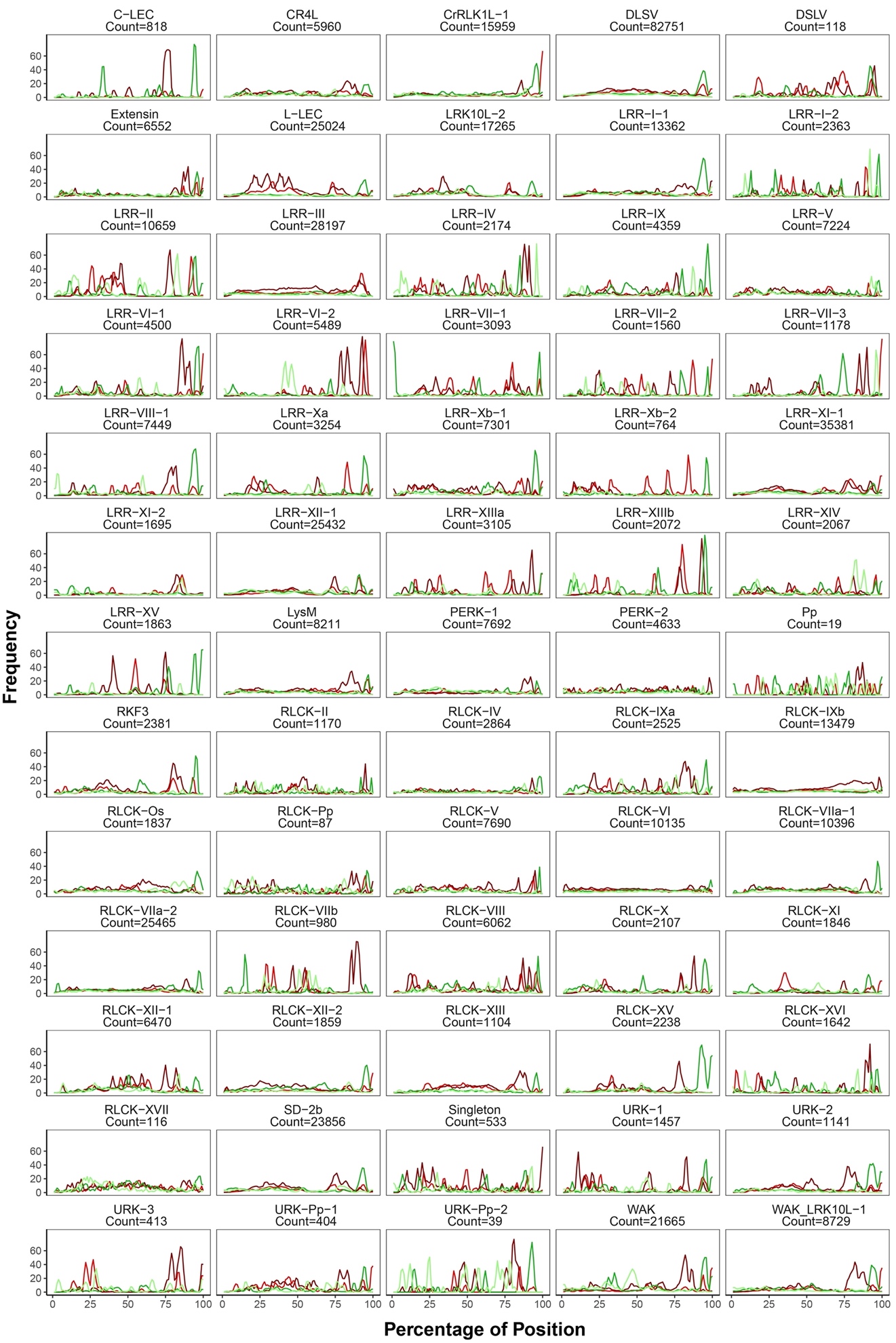

**Figure S2. Composition distribution of EDQN residues of different classes of plant RLK superfamily according to Shiu classification scheme and designation (65 classes).**

Red line refers to Aspartic acid (D), dark red line refers to Glutamic acid (E), green line refers to Asparagine (N), light green line refers to Glutamine (Q).

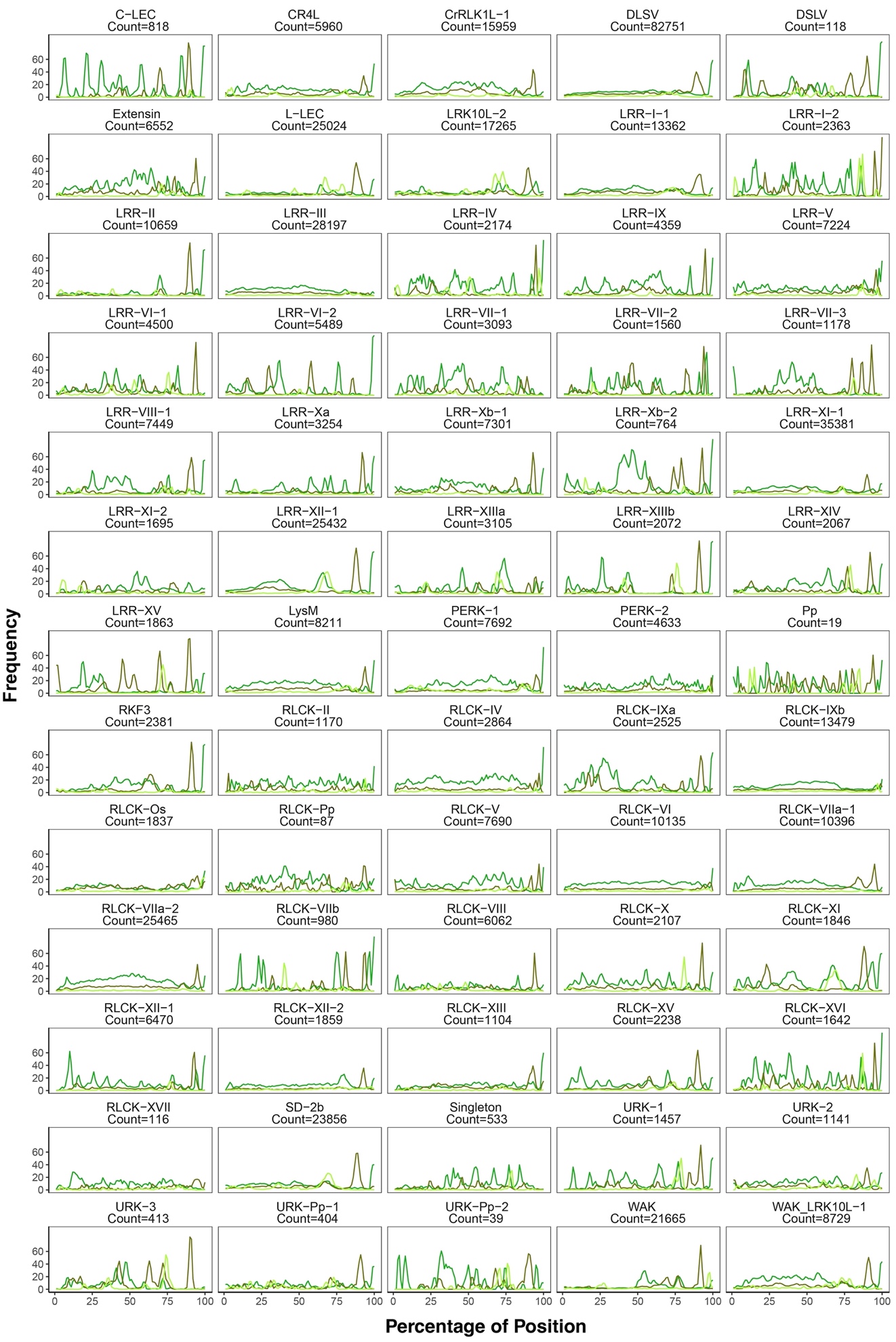

**Figure S3. Composition distribution of STY residues of different classes of plant RLK superfamily according to Shiu classification scheme and designation (65 classes).**

Green refers to Serine (S), olive refers to Threonine (T), lime refers to Tyrosine (Y).

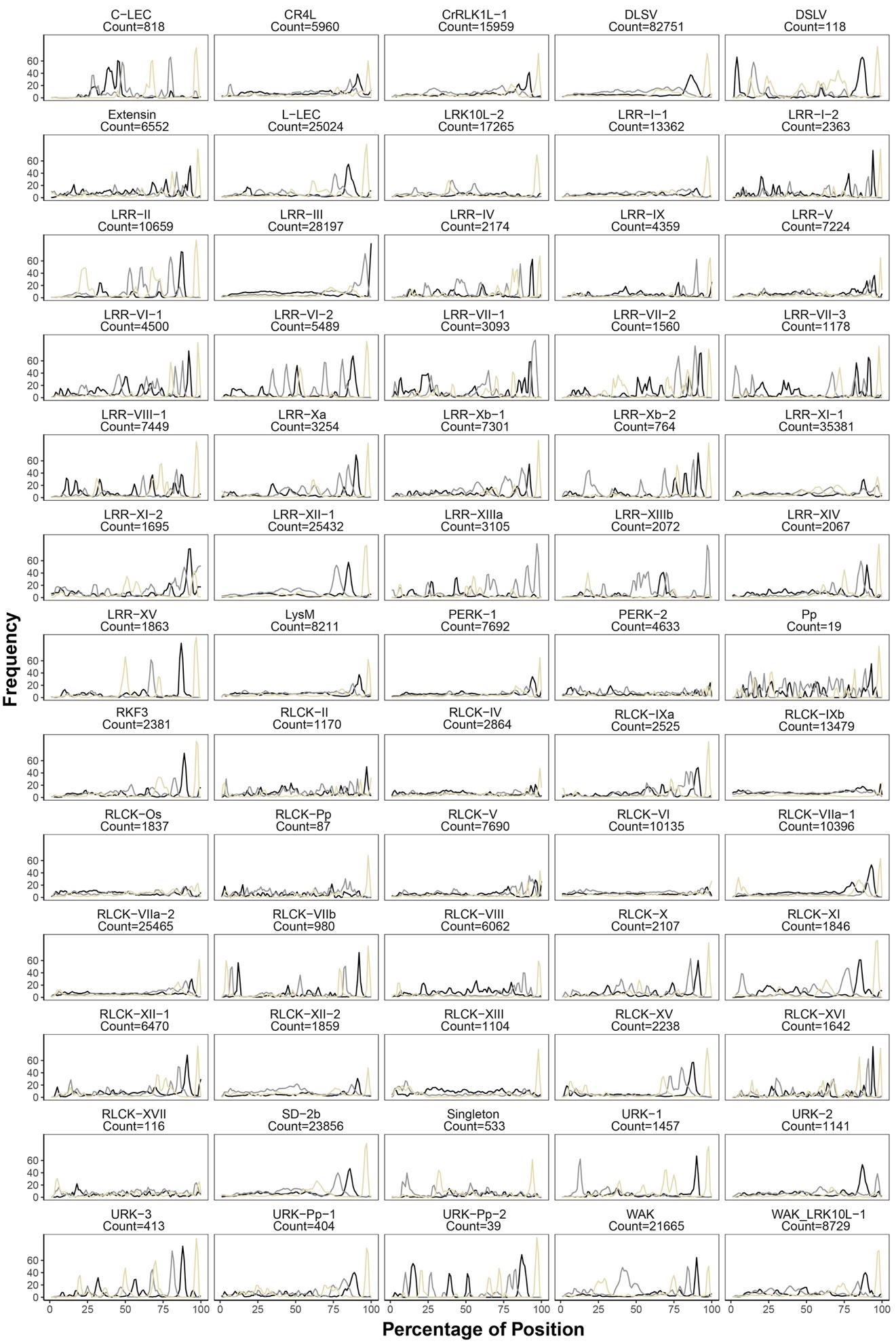

**Figure S4.** **Composition distribution of ALF residues of different classes of plant RLK superfamily according to Shiu classification scheme and designation (65 classes).**

Dark line refers to Alanine (A), grey refers to Leucine (L), beige refers to Phenylalanine (F).

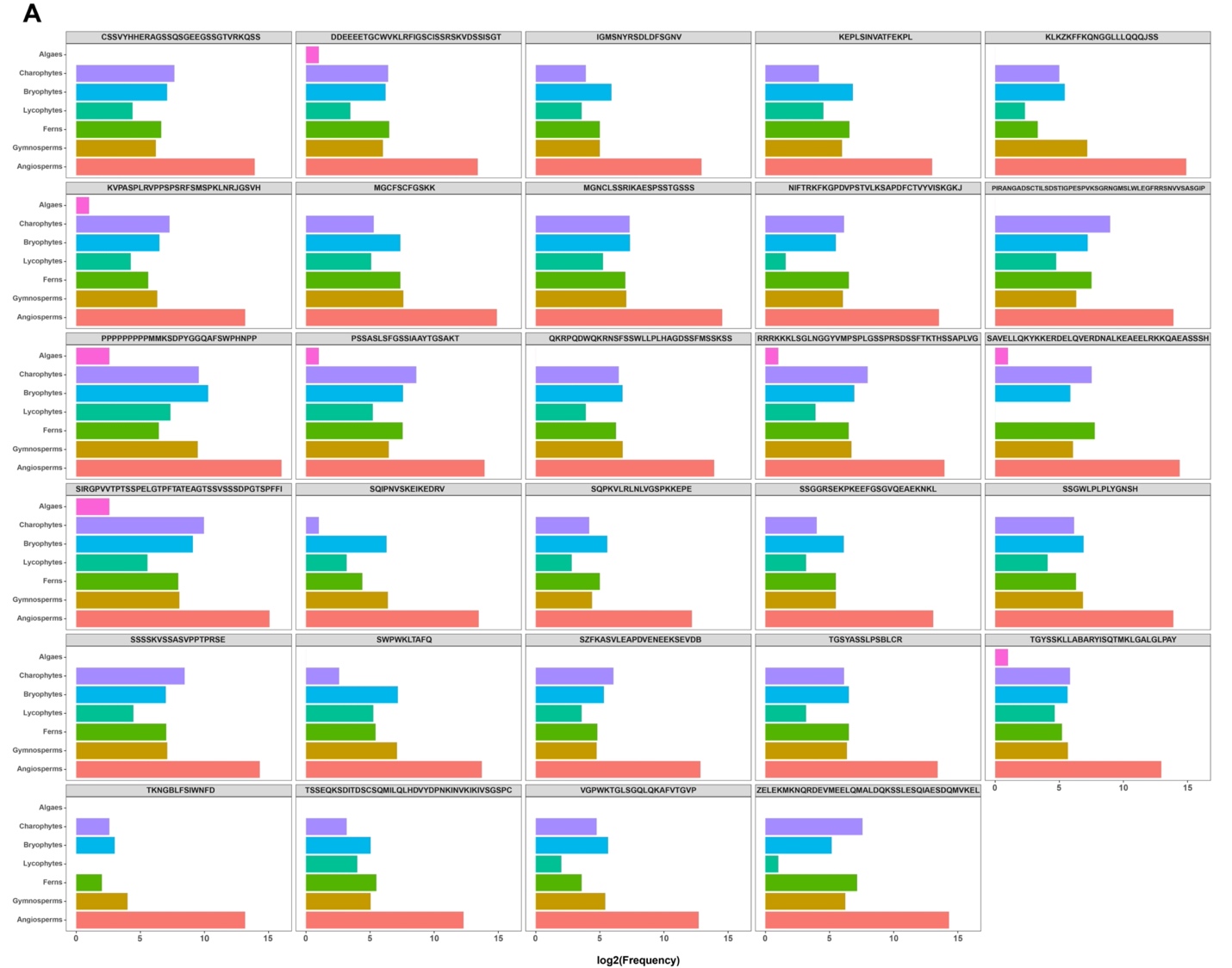

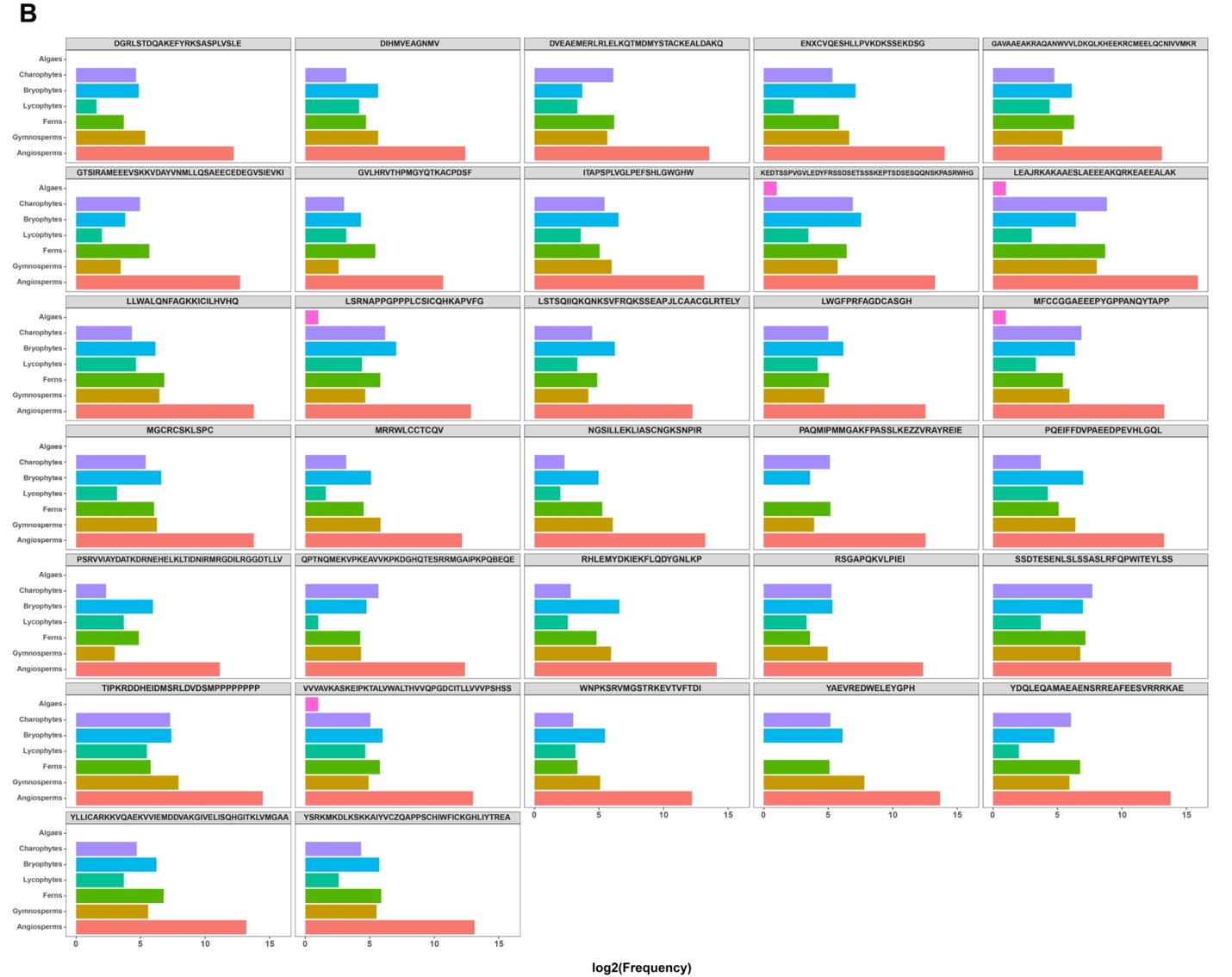

**Figure S5. The distribution of 61 motifs in 7 types of plants groups** **(Mean Similarity > 50%).**

(**A**) The distribution of 29 motifs with experimentally determined phosphorylation sites.

(**B**) The distribution of 32 motifs without experimentally determined phosphorylation sites.

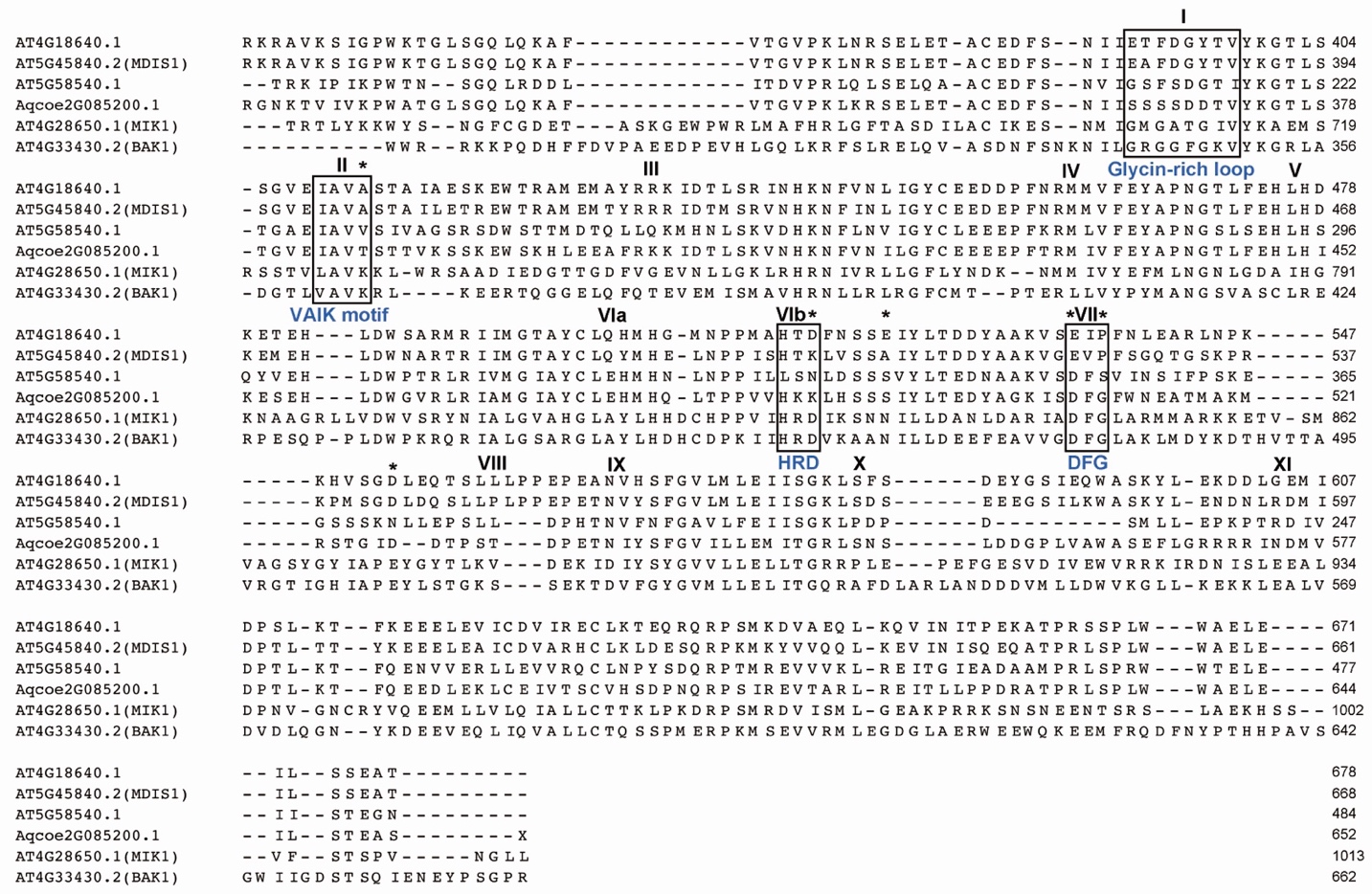

**Figure S6. Multiple sequence alignment of the kinase domain in the four representatives of the LRR-VI-2 class and the MIK1 and BAK1 RLKs.**

Roman numbers mark different subunits of the kinase domain. Asterisks on the top designate the residues conserved in active kinases. The four boxes I, II, VIb, and VII highlight critical catalytic residues (glycin-rich loop, VAIK motif, HRD and DFG motif) that LRR-VI-2 class lacks.

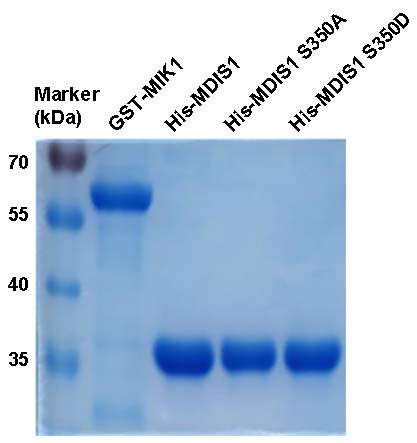

**Figure S7. The images of MIK1 and MDIS1 fusion protein gels.**

Four fusion proteins are expressed and purified before loaded for SDS-PAGE and stained with Coomassie brilliant blue G-250.

**Table S1. Source of the 528 plant species considered in the current study**

| **Class** | **Plant Species** | **Number of NRE** | **Reference** |
| --- | --- | --- | --- |
| Algaes | *Amoebophrya species* A25 | 3 | gdb database |
| Algaes | *Auxenochlorella protothecoides* | 2 | 10.1038/srep14465 |
| Algaes | *Caulerpa lentillifera* | 3 | 10.1093/dnares/dsz002 |
| Algaes | *Chlamydomonas reinhardtii* | 4 | 10.1126/science.1143609 |
| Algaes | *Chlorella variabilis* | 1 | 10.1105/tpc.110.076406 |
| Algaes | *Chromochloris zofingiensis* | 4 | 10.1073/pnas.1619928114 |
| Algaes | *Cyanophora paradoxa* | 3 | 10.1126/science.1213561 |
| Algaes | *Dunaliella salina* | 2 | 10.1128/genomea.01105-17 |
| Algaes | *Durusdinium* sp | 8 | 10.1093/gbe/evaa235 |
| Algaes | *Ectocarpus siliculosus* | 5 | 10.1111/j.1469-8137.2010.03273.x |
| Algaes | *Micromonas commoda* | 1 | 10.1126/science.1167222 |
| Algaes | *Micromonas pusilla* CCMP1545 | 1 | 10.1126/science.1167222 |
| Algaes | *Monoraphidium neglectum* | 3 | 10.1186/1471-2164-14-926 |
| Algaes | *Nemacystus decipiens* | 6 | 10.1038/s41598-019-40955-2 |
| Algaes | *Prasinoderma coloniale* | 1 | 10.1038/s41559-020-1221-7 |
| Algaes | *Pyropia yezoensis* | 1 | 10.1371/journal.pone.0057122 |
| Algaes | *Saccharina japonica* | 5 | 10.3389/fgene.2019.00378 |
| Algaes | *Ulva mutabilis* | 1 | 10.1016/j.cub.2018.08.015 |
| Algaes | *Volvox carteri* f. *nagariensis* | 2 | 10.1126/science.1188800 |
| Algaes | *Coccomyxa subellipsoidea* | 2 | 10.1186/gb-2012-13-5-r39 |
| Charophytes | *Chara braunii* | 370 | 10.1007/s12551-019-00512-7 |
| Charophytes | *Chlorokybus atmophyticus* | 11 | 10.1038/s41477-019-0560-3 |
| Charophytes | *Klebsormidium nitens* | 107 | 10.1038/ncomms4978 |
| Charophytes | *Mesostigma viride* | 5 | 10.1002/advs.201901850 |
| Charophytes | *Mesotaenium endlicherianum* | 65 | 10.1016/j.cell.2019.10.019 |
| Charophytes | *Penium margaritaceum* | 366 | 10.1016/j.cell.2020.04.019 |
| Charophytes | *Spirogloea muscicola* | 119 | 10.1016/j.cell.2019.10.019 |
| Bryophytes | *Anthoceros agrestis* | 387 | 10.1038/s41477-020-0618-2 |
| Bryophytes | *Anthoceros angustus* | 145 | 10.1038/s41477-019-0588-4 |
| Bryophytes | *Anthoceros punctatus* | 354 | 10.1038/s41477-020-0618-2 |
| Bryophytes | *Insomniella plumaeforme* | 153 | GWH database |
| Bryophytes | *Marchantia polymorpha* | 294 | 10.1016/j.cell.2017.09.030 |
| Bryophytes | *Physcomitrella patens* | 1198 | 10.1111/tpj.13801 |
| Bryophytes | *Pleurozium schreberi* | 207 | 10.1534/g3.119.400279 |
| Bryophytes | *Sphagnum fallax* | 1007 | JGI database |
| Bryophytes | *Sphagnum magellanicum* | 1002 | JGI database |
| Lycophytes | *Selaginella moellendorffii* | 771 | 10.1126/science.1203810 |
| Ferns | *Azolla filiculoides* | 318 | 10.1038/s41477-018-0188-8 |
| Ferns | *Ceratopteris richardii* | 1538 | 10.1038/s41598-019-53968-8 |
| Ferns | *Salvinia cucullata* | 285 | 10.1038/s41477-018-0188-8 |
| Gymnosperms | *Abies alba* | 149 | 10.1534/g3.119.400083 |
| Gymnosperms | *Ginkgo biloba* | 918 | 10.1038/s41477-021-00935-9 |
| Gymnosperms | *Gnetum montanum* | 350 | 10.1038/s41477-017-0097-2 |
| Gymnosperms | *Sequoiadendron giganteum* | 1014 | 10.1534/g3.120.401612 |
| Gymnosperms | *Thuja plicata* | 1953 | JGI database |
| Gymnosperms | *Welwitschia mirabilis* | 301 | 10.1038/s41467-021-24528-4 |
| Angiosperms | *Abrus precatorius* | 1036 | 10.3390/toxins11120691 |
| Angiosperms | *Acanthochlamys bracteata* | 459 | 10.1093/gbe/evab147 |
| Angiosperms | *Acer truncatum* | 422 | 10.1111/tpj.14954 |
| Angiosperms | *Acer yangbiense* | 296 | 10.1093/gigascience/giz085 |
| Angiosperms | *Acorus americanus* | 1181 | JGI database |
| Angiosperms | *Actinidia chinensis* | 861 | 10.1186/s12864-018-4656-3 |
| Angiosperms | *Actinidia eriantha* | 580 | 10.1093/gigascience/giz027 |
| Angiosperms | *Aegiceras corniculatum* | 686 | 10.1111/1755-0998.13347 |
| Angiosperms | *Aegilops tauschii* | 6296 | 10.1038/nature24486 |
| Angiosperms | *Aeschynomene evenia* | 631 | 10.1038/s41467-021-21094-7 |
| Angiosperms | *Allium sativum* | 707 | 10.1016/j.molp.2020.07.019 |
| Angiosperms | *Alnus glutinosa* | 892 | 10.5524/101042 |
| Angiosperms | *Alyssum linifolium* | 1333 | JGI database |
| Angiosperms | *Amaranthus cruentus* | 466 | 10.1111/tpj.15298 |
| Angiosperms | *Amaranthus hypochondriacus* | 334 | 10.3835/plantgenome2015.07.0062 |
| Angiosperms | *Amborella trichopoda* | 572 | 10.1126/science.1241089 |
| Angiosperms | *Ammopiptanthus nanus* | 597 | 10.5524/100466 |
| Angiosperms | *Amphicarpaea edgeworthii* | 268 | 10.1111/pbi.13520 |
| Angiosperms | *Anacardium occidentale* | 1875 | JGI database |
| Angiosperms | *Ananas bracteatus* | 446 | 10.1038/s41588-019-0506-8 |
| Angiosperms | *Ananas comosus* | 817 | 10.1038/ng.3435 |
| Angiosperms | *Antirrhinum majus* | 798 | 10.1038/s41477-018-0349-9 |
| Angiosperms | *Apium graveolens* | 446 | 10.1038/s41438-019-0235-2 |
| Angiosperms | *Aquilaria sinensis* | 581 | 10.5524/100702 |
| Angiosperms | *Aquilegia coerulea* | 859 | 10.7554/elife.36426 |
| Angiosperms | *Arabidopsis halleri* | 683 | 10.1111/1755-0998.12604 |
| Angiosperms | *Arabidopsis lyrata* | 847 | 10.1038/ng.807 |
| Angiosperms | *Arabidopsis thaliana* | 1133 | 10.1111/tpj.13415 |
| Angiosperms | *Arabis alpina* | 695 | 10.1038/nplants.2014.23 |
| Angiosperms | *Arachis duranensis* | 1210 | 10.1038/ng.3517 |
| Angiosperms | *Arachis hypogaea* | 1829 | 10.1038/s41588-019-0405-z |
| Angiosperms | *Arachis ipaensis* | 1266 | 10.1038/ng.3517 |
| Angiosperms | *Arachis monticola* | 1165 | 10.1093/gigascience/giy066 |
| Angiosperms | *Areca catechu* | 494 | 10.1111/1755-0998.13446 |
| Angiosperms | *Aristolochia fimbriata* | 370 | GWH database |
| Angiosperms | *Artocarpus altilis* | 771 | 10.3390/genes11010027 |
| Angiosperms | *Artocarpus camansi* | 685 | 10.3732/apps.1600017 |
| Angiosperms | *Artocarpus heterophyllus* | 787 | 10.3390/genes11010027 |
| Angiosperms | *Asclepias syriaca* | 287 | 10.7717/peerj.7649 |
| Angiosperms | *Asparagus officinalis* | 656 | 10.1038/s41467-017-01064-8 |
| Angiosperms | *Asparagus setaceus* | 414 | 10.5061/dryad.1c59zw3rm |
| Angiosperms | *Atalantia buxifolia* | 1164 | 10.1038/ng.3839 |
| Angiosperms | *Atropa belladonna* | 747 | MPGR database |
| Angiosperms | *Averrhoa carambola* | 477 | 10.1038/s41438-020-0307-3 |
| Angiosperms | *Avicennia marina* | 602 | 10.1093/g3journal/jkaa025 |
| Angiosperms | *Begonia fuchsioides* | 774 | 10.5524/101043 |
| Angiosperms | *Benincasa hispida* | 465 | 10.1038/s41467-019-13185-3 |
| Angiosperms | *Beta patula* | 449 | 10.1111/tpj.14413 |
| Angiosperms | *Beta vulgaris* | 749 | 10.1101/2020.09.15.298315 |
| Angiosperms | *Betula pendula* | 934 | 10.1038/ng.3862 |
| Angiosperms | *Betula platyphylla* | 998 | 10.1038/s41438-021-00481-7 |
| Angiosperms | *Boechera stricta* | 683 | 10.1038/s41559-017-0119 |
| Angiosperms | *Bombax ceiba* | 922 | 10.5524/100445 |
| Angiosperms | *Bonia amplexicaulis* | 1040 | 10.1016/j.molp.2019.05.009 |
| Angiosperms | *Botryococcus braunii* | 9 | 10.1128/genomea.00215-17 |
| Angiosperms | *Brachypodium distachyon* | 1421 | 10.1038/nature08747 |
| Angiosperms | *Brachypodium hybridum* | 1968 | 10.1038/s41467-020-17302-5 |
| Angiosperms | *Brachypodium mexicanum* | 3061 | 10.1038/s41467-020-17302-5 |
| Angiosperms | *Brachypodium stacei* | 1000 | 10.1038/s41467-020-17302-5 |
| Angiosperms | *Brachypodium sylvaticum* | 1455 | JGI database |
| Angiosperms | *Brassica chinensis* | 943 | 10.1038/s41438-020-00449-z |
| Angiosperms | *Brassica juncea* | 1370 | 10.1038/ng.3657 |
| Angiosperms | *Brassica napus* | 1906 | 10.1126/science.1253435 |
| Angiosperms | *Brassica nigra* | 937 | 10.1038/ng.3657 |
| Angiosperms | *Brassica oleracea* | 722 | 10.1186/gb-2014-15-6-r77 |
| Angiosperms | *Brassica oleracea capitata* | 534 | 10.1038/ncomms4930 |
| Angiosperms | *Brassica oleracea loquat* | 1167 | GWH database |
| Angiosperms | *Brassica rapa* | 646 | 10.1038/s41438-018-0071-9 |
| Angiosperms | *Cajanus cajan* | 684 | 10.1038/nbt.2022 |
| Angiosperms | *Cakile maritima* | 1009 | JGI database |
| Angiosperms | *Calamus simplicifolius* | 904 | 10.5524/101052 |
| Angiosperms | *Calotropis gigantea* | 417 | 10.1534/g3.117.300331 |
| Angiosperms | *Camelina sativa* | 1810 | 10.1038/ncomms4706 |
| Angiosperms | *Camellia sinensis* | 1588 | 10.1016/j.molp.2020.04.010 |
| Angiosperms | *Camptotheca acuminata* | 1433 | 10.1038/s41467-021-23872-9 |
| Angiosperms | *Cannabis sativa* | 993 | 10.1038/s41438-020-0295-3 |
| Angiosperms | *Capsella grandiflora* | 626 | 10.1038/ng.2669 |
| Angiosperms | *Capsella rubella* | 878 | 10.1038/ng.2669 |
| Angiosperms | *Capsicum annuum* | 837 | 10.1073/pnas.1400975111 |
| Angiosperms | *Capsicum baccatum* | 745 | 10.1186/s13059-017-1341-9 |
| Angiosperms | *Capsicum chinense* | 735 | 10.1186/s13059-017-1341-9 |
| Angiosperms | *Cardamine hirsuta* | 855 | 10.1038/nplants.2016.167 |
| Angiosperms | *Carica papaya* | 511 | 10.1038/nature06856 |
| Angiosperms | *Carthamus tinctorius* | 726 | 10.1111/pbi.13586 |
| Angiosperms | *Carya cathayensis* | 756 | 10.1093/gigascience/giz036 |
| Angiosperms | *Carya illinoinensis* | 2332 | 10.1038/s41467-021-24328-w |
| Angiosperms | *Castanea dentata* | 1787 | JGI database |
| Angiosperms | *Castanea mollissima* | 992 | 10.1038/nature20786 |
| Angiosperms | *Casuarina glauca* | 599 | 10.5524/101051 |
| Angiosperms | *Catharanthus roseus* | 883 | 10.5061/dryad.hs593 |
| Angiosperms | *Caulanthus amplexicaulis* | 985 | JGI database |
| Angiosperms | *Cenchrus americanus* | 741 | 10.1038/nbt.3943 |
| Angiosperms | *Cenchrus purpureus* | 1307 | 10.1111/1755-0998.13271 |
| Angiosperms | *Cephalotus follicularis* | 352 | 10.5061/dryad.50tq3 |
| Angiosperms | *Cerasus yedoensis* | 1693 | 10.1186/s13059-018-1497-y |
| Angiosperms | *Ceratodon purpureus* GG1 | 606 | 10.1126/sciadv.abh2488 |
| Angiosperms | *Ceratodon purpureus* R40 | 583 | 10.1126/sciadv.abh2488 |
| Angiosperms | *Cercis canadensis* | 694 | 10.5524/101045 |
| Angiosperms | *Chamaecrista fasciculata* | 655 | 10.5524/101045 |
| Angiosperms | *Chenopodium pallidicaule* | 278 | 10.1002/aps3.11300 |
| Angiosperms | *Chenopodium quinoa* | 1217 | 10.1038/nature21370 |
| Angiosperms | *Chenopodium suecicum* | 349 | 10.1038/nature21370 |
| Angiosperms | *Chimonanthus praecox* | 556 | 10.1186/s13059-020-02088-y |
| Angiosperms | *Chimonanthus salicifolius* | 484 | 10.1111/tpj.14874 |
| Angiosperms | *Chiococca alba* | 1053 | 10.5061/dryad.00000000r |
| Angiosperms | *Chrysanthemum nankingense* | 1970 | 10.1016/j.molp.2018.10.003 |
| Angiosperms | *Chrysanthemum seticuspe* | 2145 | 10.1093/dnares/dsy048 |
| Angiosperms | *Cicer arietinum* | 891 | 10.1038/nbt.2491 |
| Angiosperms | *Cinnamomum kanehirae* | 919 | 10.1038/s41477-018-0337-0 |
| Angiosperms | *Cinnamomum micranthum* | 885 | 10.1038/s41477-018-0337-0 |
| Angiosperms | *Citrullus lanatus* | 467 | 10.1111/pbi.13136 |
| Angiosperms | *Citrus clementina* | 904 | 10.1038/nature25447 |
| Angiosperms | *Citrus grandis* | 951 | 10.1038/ng.3839 |
| Angiosperms | *Citrus ichangensis* | 564 | 10.1038/ng.3839 |
| Angiosperms | *Citrus medica* | 520 | 10.1038/ng.3839 |
| Angiosperms | *Citrus reticulata* | 1016 | 10.1016/j.molp.2018.06.001 |
| Angiosperms | *Citrus sinensis* | 1160 | 10.1038/nbt.2906 |
| Angiosperms | *Citrus unshiu* | 1071 | 10.3389/fgene.2017.00180 |
| Angiosperms | *Cladopus chinensis* | 386 | 10.1038/s41438-020-0269-5 |
| Angiosperms | *Cleistogenes songorica* | 1095 | 10.1111/pbi.13483 |
| Angiosperms | *Cleome violacea* | 616 | JGI database |
| Angiosperms | *Cocos nucifera* | 616 | 10.5524/100347 |
| Angiosperms | *Coffea arabica* | 1835 | 10.1111/pbi.12912 |
| Angiosperms | *Coffea canephora* | 784 | 10.1126/science.1255274 |
| Angiosperms | *Coffea eugenioides* | 1157 | NCBI database |
| Angiosperms | *Coffea humblotiana* | 444 | 10.1038/s41598-021-87419-0 |
| Angiosperms | *Coix lacryma-jobi* | 830 | 10.3389/fpls.2020.00630 |
| Angiosperms | *Corchorus capsularis* | 355 | 10.1038/nplants.2016.223 |
| Angiosperms | *Corchorus olitorius* | 525 | 10.1016/j.gdata.2017.05.007 |
| Angiosperms | *Coriandrum sativum* | 798 | 10.1111/pbi.13310 |
| Angiosperms | *Corylus avellana* | 588 | 10.1111/tpj.15099 |
| Angiosperms | *Corymbia citriodora* | 2055 | 10.1038/s42003-021-02009-0 |
| Angiosperms | *Crambe hispanica* | 945 | JGI database |
| Angiosperms | *Crucihimalaya himalaica* | 471 | 10.1073/pnas.1817580116 |
| Angiosperms | *Cucumis hystrix* | 354 | 10.1038/s41438-021-00475-5 |
| Angiosperms | *Cucumis melo* | 648 | 10.1073/pnas.1205415109 |
| Angiosperms | *Cucumis sativus* | 722 | 10.1093/gigascience/giz072 |
| Angiosperms | *Cucumis x hytivus* | 604 | 10.1002/advs.202004222 |
| Angiosperms | *Cucurbita argyrosperma* | 524 | 10.1038/s41438-021-00544-9 |
| Angiosperms | *Cucurbita maxima* | 1033 | 10.1016/j.molp.2017.09.003 |
| Angiosperms | *Cucurbita moschata* | 1072 | 10.1016/j.molp.2017.09.003 |
| Angiosperms | *Cucurbita pepo* | 1033 | 10.1111/pbi.12860 |
| Angiosperms | *Cuscuta australis* | 314 | 10.1038/s41467-018-04721-8 |
| Angiosperms | *Cuscuta campestris* | 744 | 10.1038/s41467-018-04344-z |
| Angiosperms | *Cynara cardunculus* | 838 | 10.1038/s41598-017-05085-7 |
| Angiosperms | *Cynara cardunculus* var. *scolymus* | 495 | 10.1038/s41598-017-05085-7 |
| Angiosperms | *Daemonorops jenkinsiana* | 1215 | 10.5524/101053 |
| Angiosperms | *Datisca glomerata* | 479 | 10.5524/101046 |
| Angiosperms | *Daucus carota* | 1378 | 10.1038/ng.3565 |
| Angiosperms | *Davidia involucrata* | 940 | 10.1111/1755-0998.13138 |
| Angiosperms | *Dendrobium catenatum* | 696 | 10.1038/srep19029 |
| Angiosperms | *Dendrobium huoshanense* | 323 | 10.1093/gbe/evaa215 |
| Angiosperms | *Dendrobium officinale* | 719 | 10.1016/j.apsb.2021.01.019 |
| Angiosperms | *Descurainia sophioides* | 729 | JGI database |
| Angiosperms | *Dianthus caryophyllus* | 829 | 10.1093/dnares/dst053 |
| Angiosperms | *Digitalis purpurea* | 1374 | MPGR database |
| Angiosperms | *Digitaria exilis* | 1652 | 10.1093/gigascience/giab013 |
| Angiosperms | *Dimocarpus longan* Lour | 967 | 10.5524/100276 |
| Angiosperms | *Dioscorea alata* | 970 | 10.1101/2021.04.14.439117 |
| Angiosperms | *Dioscorea cayenensis* | 835 | 10.1111/mpp.12137 |
| Angiosperms | *Dioscorea dumetorum* | 697 | 10.3390/genes11030274 |
| Angiosperms | *Dioscorea rotundata* | 1179 | 10.1186/s12915-017-0419-x |
| Angiosperms | *Dioscorea villosa* | 621 | MPGR database |
| Angiosperms | *Diospyros lotus* | 866 | 10.1371/journal.pgen.1008566 |
| Angiosperms | *Diospyros oleifera* | 539 | 10.1038/s41438-019-0227-2 |
| Angiosperms | *Diptychocarpus strictus* | 827 | JGI database |
| Angiosperms | *Discaria trinervis* | 539 | 10.5524/101048 |
| Angiosperms | *Draba nivalis* | 596 | 10.1111/1755-0998.13280 |
| Angiosperms | *Dryas drummondii* | 585 | 10.5524/101047 |
| Angiosperms | *Durio zibethinus* | 1656 | 10.1038/ng.3972 |
| Angiosperms | *Echinacea purpurea* | 1282 | MPGR database |
| Angiosperms | *Echinochloa crus-galli* | 1765 | 10.1016/j.molp.2020.07.001 |
| Angiosperms | *Elaeis guineensis* | 1166 | 10.1038/nature12309 |
| Angiosperms | *Eleusine coracana* | 1321 | 10.1186/s12864-017-3850-z |
| Angiosperms | *Ensete glaucum* | 719 | 10.1093/database/bat035 |
| Angiosperms | *Ensete ventricosum* | 477 | 10.1016/j.dib.2018.03.026 |
| Angiosperms | *Eragrostis curvula* | 1052 | 10.1038/s41598-019-46610-0 |
| Angiosperms | *Eragrostis tef* | 1217 | 10.1186/1471-2164-15-581 |
| Angiosperms | *Erigeron breviscapus* | 529 | 10.5524/100290 |
| Angiosperms | *Eriobotrya japonica* | 1183 | 10.5524/100711 |
| Angiosperms | *Eruca vesicaria* | 1705 | JGI database |
| Angiosperms | *Erysimum cheiranthoides* | 574 | 10.7554/eLife.51712 |
| Angiosperms | *Eschscholzia californica* | 616 | 10.1093/bbb/zbaa091 |
| Angiosperms | *Eucalyptus camaldulensis* | 1622 | 10.5511/plantbiotechnology.11.1027b |
| Angiosperms | *Eucalyptus grandis* | 2164 | 10.1111/nph.13150 |
| Angiosperms | *Euclidium syriacum* | 815 | JGI database |
| Angiosperms | *Eucommia ulmoides* | 299 | 10.1038/s41438-020-00406-w |
| Angiosperms | *Eutrema salsugineum* | 780 | 10.3389/fpls.2013.00046 |
| Angiosperms | *Fagopyrum tataricum* | 705 | 10.1016/j.molp.2017.08.013 |
| Angiosperms | *Fagus sylvatica* | 1307 | 10.1093/gigascience/giy063 |
| Angiosperms | *Faidherbia albida* | 681 | 10.5524/101054 |
| Angiosperms | *Ficus carica* | 613 | 10.1038/srep41124 |
| Angiosperms | *Ficus erecta* | 1426 | 10.1111/tpj.14703 |
| Angiosperms | *Ficus hispida* | 672 | 10.1016/j.cell.2020.09.043 |
| Angiosperms | *Ficus microcarpa* | 747 | 10.1016/j.cell.2020.09.043 |
| Angiosperms | *Fortunella hindsii* | 871 | 10.1111/pbi.13132 |
| Angiosperms | *Fragaria ananassa* | 2056 | 10.1038/s41588-019-0356-4 |
| Angiosperms | *Fragaria iinumae* | 2056 | 10.1038/s41588-019-0544-2 |
| Angiosperms | *Fragaria nilgerrensis* | 641 | 10.1111/pbi.13351 |
| Angiosperms | *Fragaria vesca* | 1493 | 10.1093/gigascience/gix124 |
| Angiosperms | *Fraxinus excelsior* | 1214 | 10.1038/nature20786 |
| Angiosperms | *Gastrodia elata* | 301 | 10.1038/s41467-018-03423-5 |
| Angiosperms | *Gillenia trifoliata* | 549 | 10.1038/s41438-021-00662-4 |
| Angiosperms | *Glycine max* | 2174 | 10.1038/nature08670 |
| Angiosperms | *Glycine soja* | 2423 | 10.1111/tpj.14500 |
| Angiosperms | *Gossypioides kirkii* | 657 | 10.3389/fpls.2019.01541 |
| Angiosperms | *Gossypium arboreum* | 1236 | 10.1038/ng.2987 |
| Angiosperms | *Gossypium aridum* | 576 | 10.1038/s41588-020-0607-4 |
| Angiosperms | *Gossypium armourianum* | 624 | 10.1093/gbe/evy256 |
| Angiosperms | *Gossypium australe* | 654 | 10.1111/pbi.13249 |
| Angiosperms | *Gossypium barbadense* | 2638 | 10.1038/s41588-020-0614-5 |
| Angiosperms | *Gossypium darwinii* | 2340 | 10.1038/s41588-020-0614-5 |
| Angiosperms | *Gossypium davidsonii* | 986 | 10.1186/s12915-021-01041-0 |
| Angiosperms | *Gossypium gossypioides* | 585 | 10.1093/gbe/evy256 |
| Angiosperms | *Gossypium harknessii* | 643 | 10.1093/gbe/evy256 |
| Angiosperms | *Gossypium herbaceum* | 771 | 10.1038/s41588-020-0607-4 |
| Angiosperms | *Gossypium hirsutum* | 2507 | 10.1038/s41588-020-0614-5 |
| Angiosperms | *Gossypium lobatum* | 624 | 10.1093/gbe/evy256 |
| Angiosperms | *Gossypium longicalyx* | 914 | 10.1534/g3.120.401050 |
| Angiosperms | *Gossypium mustelinum* | 2485 | 10.1038/s41588-020-0614-5 |
| Angiosperms | *Gossypium raimondii* | 1645 | 10.1038/nature11798 |
| Angiosperms | *Gossypium tomentosum* | 2597 | 10.1038/s41588-020-0614-5 |
| Angiosperms | *Guadua angustifolia* | 1224 | 10.1016/j.molp.2019.05.009 |
| Angiosperms | *Handroanthus impetiginosus* | 957 | 10.1093/gigascience/gix125 |
| Angiosperms | *Helianthus annuus* | 2074 | 10.1038/nature22380 |
| Angiosperms | *Herrania umbratica* | 780 | NCBI database |
| Angiosperms | *Hevea brasiliensis* | 1818 | 10.1038/nplants.2016.73 |
| Angiosperms | *Hibiscus cannabinus* | 143 | 10.1111/pbi.13341 |
| Angiosperms | *Hibiscus syriacus* | 1853 | 10.1093/dnares/dsw049 |
| Angiosperms | *Hoodia gordonii* | 181 | MPGR database |
| Angiosperms | *Hordeum vulgare* | 6278 | 10.1038/sdata.2017.44 |
| Angiosperms | *Hydrangea macrophylla* | 844 | 10.1101/2020.06.14.151431 |
| Angiosperms | *Hydrangea quercifolia* | 1272 | JGI database |
| Angiosperms | *Hypericum perforatum* | 944 | 10.1111/jpi.12709 |
| Angiosperms | *Iberis amara* | 1060 | JGI database |
| Angiosperms | *Ipomoea batatas* | 885 | 10.1007/s00299-019-02464-4 |
| Angiosperms | *Ipomoea nil* | 1431 | 10.1038/ncomms13295 |
| Angiosperms | *Ipomoea trifida* | 1228 | 10.1038/s41467-018-06983-8 |
| Angiosperms | *Ipomoea triloba* | 1372 | 10.1038/s41467-018-06983-8 |
| Angiosperms | *Isatis tinctoria* | 2269 | JGI database |
| Angiosperms | *Jacaranda mimosifolia* | 526 | 10.1093/gbe/evab094 |
| Angiosperms | *Jatropha curcas* | 820 | 10.1093/dnares/dsq030 |
| Angiosperms | *Joinvillea ascendens* | 1170 | JGI database |
| Angiosperms | *Juglans regia* | 1516 | 10.1093/gigascience/giaa050 |
| Angiosperms | *Kalanchoe fedtschenkoi* | 875 | 10.1038/s41467-017-01491-7 |
| Angiosperms | *Kalanchoe laxiflora* | 1528 | JGI database |
| Angiosperms | *Kandelia obovata* | 335 | 10.1038/s41438-020-0300-x |
| Angiosperms | *Lablab purpureus* | 410 | 10.5524/101056 |
| Angiosperms | *Lactuca sativa* | 1846 | 10.1038/ncomms14953 |
| Angiosperms | *Lagenaria siceraria* | 475 | 10.1111/tpj.13722 |
| Angiosperms | *Leersia perrieri* | 1102 | 10.1038/s41588-018-0040-0 |
| Angiosperms | *Lepidium sativum* | 1176 | JGI database |
| Angiosperms | *Lindenbergia philippensis* | 1138 | JGI database |
| Angiosperms | *Liriodendron chinense* | 921 | 10.1038/s41477-018-0323-6 |
| Angiosperms | *Lobularia maritima* | 562 | 10.1038/s41438-020-00422-w |
| Angiosperms | *Lonicera japonica* | 649 | 10.1111/nph.16552 |
| Angiosperms | *Lotus japonicus* | 803 | 10.3390/genes11050483 |
| Angiosperms | *Luffa cylindrica* | 524 | 10.1111/1755-0998.13129 |
| Angiosperms | *Lunaria annua* | 831 | JGI database |
| Angiosperms | *Lupinus albus* | 698 | 10.1038/s41467-019-14197-9 |
| Angiosperms | *Lupinus angustifolius* | 1217 | 10.1111/pbi.12615 |
| Angiosperms | *Macadamia integrifolia* | 1150 | 10.1186/s12864-016-3272-3 |
| Angiosperms | *Macleaya cordata* | 347 | 10.1016/j.molp.2017.05.007 |
| Angiosperms | *Macrotyloma uniflorum* | 736 | 10.1101/2021.01.18.427074 |
| Angiosperms | *Magnolia biondii* | 572 | 10.1038/s41438-021-00471-9 |
| Angiosperms | *Malania oleifera* | 464 | 10.5524/100549 |
| Angiosperms | *Malcolmia maritima* | 696 | JGI database |
| Angiosperms | *Malus baccata* | 879 | 10.1534/g3.119.400245 |
| Angiosperms | *Malus domestica* | 1578 | 10.1038/ng.3886 |
| Angiosperms | *Mangifera indica* | 535 | 10.1186/s13059-020-01959-8 |
| Angiosperms | *Manihot esculenta* | 1829 | Bredeson et al.,In preparation. |
| Angiosperms | *Medicago polymorpha* | 701 | 10.1038/s41438-021-00483-5 |
| Angiosperms | *Medicago sativa* | 3704 | 10.1038/s41467-020-16338-x |
| Angiosperms | *Medicago truncatula* | 1230 | 10.1038/nature10625 |
| Angiosperms | *Megacarpaea delavayi* | 791 | 10.3389/fgene.2020.00812 |
| Angiosperms | *Megadenia pygmaea* Mpyg | 534 | 10.1111/1755-0998.13291 |
| Angiosperms | *Mercurialis annua* | 557 | 10.1534/genetics.119.302045 |
| Angiosperms | *Mikania micrantha* | 1008 | 10.1038/s41467-019-13926-4 |
| Angiosperms | *Mimosa pudica* | 633 | 10.5524/101049 |
| Angiosperms | *Mimulus guttatus* | 837 | 10.1073/pnas.1319032110 |
| Angiosperms | *Mimulus guttatus* NONTOL | 892 | JGI database |
| Angiosperms | *Mimulus guttatus* TOL | 887 | JGI database |
| Angiosperms | *Miscanthus lutarioriparius* | 1344 | 10.1038/s41467-021-22738-4 |
| Angiosperms | *Miscanthus sinensis* | 2410 | 10.1038/s41467-020-18923-6 |
| Angiosperms | *Mitragyna speciosa* | 2064 | 10.1093/g3journal/jkab058 |
| Angiosperms | *Momordica charantia* | 671 | 10.1073/pnas.1921016117 |
| Angiosperms | *Moringa oleifera* | 358 | 10.5524/101058 |
| Angiosperms | *Morus notabilis* | 548 | 10.1038/ncomms3445 |
| Angiosperms | *Musa acuminata* | 1302 | 10.1038/nature11241 |
| Angiosperms | *Musa balbisiana* | 576 | 10.1038/s41477-019-0452-6 |
| Angiosperms | *Musa itinerans* | 513 | 10.1093/database/bat035 |
| Angiosperms | *Musa schizocarpa* | 703 | 10.1093/database/bat035 |
| Angiosperms | *Myagrum perfoliatum* | 712 | JGI database |
| Angiosperms | *Nelumbo nucifera* | 981 | 10.1111/tpj.13894 |
| Angiosperms | *Nicotiana attenuata* | 799 | 10.1073/pnas.1700073114 |
| Angiosperms | *Nicotiana benthamiana* | 877 | 10.1094/MPMI-06-12-0148-TA |
| Angiosperms | *Nicotiana sylvestris* | 1084 | 10.1186/gb-2013-14-6-r60 |
| Angiosperms | *Nicotiana tabacum* | 1584 | 10.1186/s12864-017-3791-6 |
| Angiosperms | *Nicotiana tomentosiformis* | 1047 | 10.3732/ajb.89.6.921 |
| Angiosperms | *Nissolia schottii* | 658 | 10.5524/101050 |
| Angiosperms | *Nymphaea colorata* | 523 | 10.1038/s41586-019-1852-5 |
| Angiosperms | *Nymphaea thermarum* | 481 | 10.1073/pnas.1922873117 |
| Angiosperms | *Ocimum basilicum* | 1837 | 10.1093/dnares/dsaa016 |
| Angiosperms | *Olea europaea* | 1063 | 10.1073/pnas.1708621114 |
| Angiosperms | *Olyra latifolia* | 752 | 10.1016/j.molp.2019.05.009 |
| Angiosperms | *Ophiorrhiza pumila* | 489 | 10.1038/s41467-020-20508-2 |
| Angiosperms | *Origanum majorana* | 827 | 10.1093/dnares/dsaa016 |
| Angiosperms | *Origanum vulgare* | 840 | 10.1093/dnares/dsaa016 |
| Angiosperms | *Oropetium thomaeum* | 87 | 10.1038/nature15714 |
| Angiosperms | *Oryza barthii* | 1028 | 10.1270/jsbbs.17022 |
| Angiosperms | *Oryza brachyantha* | 919 | 10.1038/ncomms2596 |
| Angiosperms | *Oryza glaberrima* | 811 | 10.1038/ng.3044 |
| Angiosperms | *Oryza glumipatula* | 1165 | ensembl database |
| Angiosperms | *Oryza granulata* | 587 | 10.1038/s41597-020-0470-2 |
| Angiosperms | *Oryza indica* | 888 | ensembl database |
| Angiosperms | *Oryza longistaminata* | 923 | 10.1002/tpg2.20001 |
| Angiosperms | *Oryza meridionalis* | 1073 | 10.1073/pnas.1418307111 |
| Angiosperms | *Oryza nivara* | 1213 | 10.1073/pnas.1418307111 |
| Angiosperms | *Oryza punctata* | 1012 | 10.1038/s41588-018-0040-0 |
| Angiosperms | *Oryza rufipogon* | 1131 | 10.1007/s11427-020-1738-x |
| Angiosperms | *Oryza sativa* | 1205 | 10.1093/nar/gkl976 |
| Angiosperms | *Ostrya chinensis* | 332 | 10.1038/s41467-018-07913-4 |
| Angiosperms | *Ostrya rehderiana* | 484 | 10.1038/s41467-018-07913-4 |
| Angiosperms | *Panax ginseng* | 1100 | 10.1038/s41597-019-0296-y |
| Angiosperms | *Panax notoginseng* | 560 | 10.1016/j.isci.2020.101538 |
| Angiosperms | *Panax quinquefolius* | 1029 | MPGR database |
| Angiosperms | *Panicum hallii* | 1118 | 10.1038/s41467-018-07669-x |
| Angiosperms | *Panicum miliaceum* | 1170 | 10.1038/s41467-019-08409-5 |
| Angiosperms | *Panicum virgatum* | 3530 | 10.1038/s41586-020-03127-1 |
| Angiosperms | *Papaver somniferum* | 1574 | 10.1126/science.aat4096 |
| Angiosperms | *Paspalum vaginatum* | 1462 | JGI database |
| Angiosperms | *Passiflora edulis* | 433 | 10.1111/1755-0998.13310 |
| Angiosperms | *Petunia axillaris* | 581 | 10.1038/nplants.2016.74 |
| Angiosperms | *Petunia inflata* | 688 | 10.1038/nplants.2016.74 |
| Angiosperms | *Phalaenopsis aphrodite* | 407 | 10.1111/pbi.12936 |
| Angiosperms | *Phalaenopsis equestris* | 585 | 10.1038/ng.3149 |
| Angiosperms | *Pharus latifolius* | 1670 | JGI database |
| Angiosperms | *Phaseolus acutifolius* | 1455 | 10.1038/s41467-021-22858-x |
| Angiosperms | *Phaseolus acutifolius* WLD | 966 | 10.1038/s41467-021-22858-x |
| Angiosperms | *Phaseolus lunatus* | 966 | 10.1038/s41467-021-20921-1 |
| Angiosperms | *Phaseolus vulgaris* | 1081 | 10.1038/ng.3008 |
| Angiosperms | *Phaseolus vulgaris* Labor Ovalle | 1423 | JGI database |
| Angiosperms | *Phoenix dactylifera* | 1071 | 10.1038/ncomms3274 |
| Angiosperms | *Phtheirospermum japonicum* | 515 | 10.1126/sciadv.abc2385 |
| Angiosperms | *Phyllostachys edulis* | 1202 | 10.5524/100498 |
| Angiosperms | *Phytolacca americana* | 1038 | 10.3389/fpls.2019.01002 |
| Angiosperms | *Piper nigrum* | 954 | 10.1038/s41467-019-12607-6 |
| Angiosperms | *Pistacia vera* | 857 | 10.1186/s12864-016-3359-x |
| Angiosperms | *Pisum sativum* | 574 | 10.1101/2020.09.25.313072 |
| Angiosperms | *Poncirus trifoliata* | 968 | 10.1111/tpj.14993 |
| Angiosperms | *Populus alba* | 1541 | 10.1111/pbi.12989 |
| Angiosperms | *Populus deltoides* WV94 | 1589 | 10.1093/jhered/esab010 |
| Angiosperms | *Populus euphratica* | 728 | 10.1111/1755-0998.13142 |
| Angiosperms | *Populus pruinosa* | 866 | 10.5524/100319 |
| Angiosperms | *Populus tremula* | 1856 | 10.1073/pnas.1801437115 |
| Angiosperms | *Populus tremuloides* | 909 | 10.1073/pnas.1801437115 |
| Angiosperms | *Populus trichocarpa* | 1780 | 10.1126/science.1128691 |
| Angiosperms | *Populus trichocarpa* Stettler14 | 1655 | 10.1534/g3.119.400913 |
| Angiosperms | *Portulaca amilis* | 859 | 10.1093/plphys/kiac116 |
| Angiosperms | *Potentilla micrantha* | 311 | 10.5524/100407 |
| Angiosperms | *Primula veris* | 433 | 10.5061/dryad.2s200 |
| Angiosperms | *Prosopis alba* | 1946 | NCBI database |
| Angiosperms | *Prunus armeniaca* | 1278 | 10.1038/s41438-019-0215-6 |
| Angiosperms | *Prunus avium* | 496 | 10.1093/dnares/dsx020 |
| Angiosperms | *Prunus domestica* | 1196 | 10.1038/s41438-020-00438-2 |
| Angiosperms | *Prunus dulcis* | 863 | 10.1111/tpj.14538 |
| Angiosperms | *Prunus mandshurica* | 948 | rosaceae database |
| Angiosperms | *Prunus mume* | 907 | 10.1038/ncomms2290 |
| Angiosperms | *Prunus persica* | 923 | 10.1038/ng.2586 |
| Angiosperms | *Prunus sibirica* | 1013 | rosaceae database |
| Angiosperms | *Puccinellia tenuiflora* | 828 | 10.1007/s11427-020-1662-x |
| Angiosperms | *Punica granatum* | 993 | 10.1111/pbi.12875 |
| Angiosperms | *Pyrus betulifolia* | 1202 | 10.1111/pbi.13226. |
| Angiosperms | *Pyrus communis* | 859 | 10.1093/gigascience/giz138 |
| Angiosperms | *Pyrus pyrifolia* | 1155 | 10.1093/dnares/dsab001 |
| Angiosperms | *Pyrus x bretschneideri* | 1359 | 10.1101/gr.144311.112 |
| Angiosperms | *Quercus lobata* | 1482 | 10.1534/g3.116.030411 |
| Angiosperms | *Quercus robur* | 978 | 10.1038/s41477-018-0172-3 |
| Angiosperms | *Quercus rubra* | 1845 | JGI database |
| Angiosperms | *Quercus suber* | 1643 | 10.1038/sdata.2018.69 |
| Angiosperms | *Raddia distichophylla* | 435 | 10.1093/g3journal/jkaa049 |
| Angiosperms | *Raddia guianensis* | 579 | 10.1016/j.molp.2019.05.009 |
| Angiosperms | *Raphanus raphanistrum* | 913 | 10.1105/tpc.114.124297 |
| Angiosperms | *Raphanus raphanistrum* subsp. *landra* | 924 | GWH database |
| Angiosperms | *Raphanus sativus* | 1081 | 10.1093/dnares/dsaa010 |
| Angiosperms | *Rauvolfia serpentina* | 1052 | MPGR database |
| Angiosperms | *Rhodamnia argentea* | 1152 | NCBI database |
| Angiosperms | *Rhodiola crenulata* | 499 | 10.5524/100301 |
| Angiosperms | *Rhododendron delavayi* | 760 | 10.5524/100331 |
| Angiosperms | *Rhododendron ovatum* | 1271 | 10.1111/pbi.13680 |
| Angiosperms | *Rhododendron ripense* | 859 | 10.1101/2021.06.17.448907 |
| Angiosperms | *Rhododendron simsii* | 710 | 10.1038/s41467-020-18771-4. |
| Angiosperms | *Rhododendron williamsianum* | 270 | 10.1093/gbe/evz245. |
| Angiosperms | *Ricinus communis* | 790 | 10.1038/nbt.1674 |
| Angiosperms | *Rorippa islandica* | 712 | JGI database |
| Angiosperms | *Rosa chinensis* | 1158 | 10.1038/s41477-018-0166-1 |
| Angiosperms | *Rosa multiflora* | 1395 | 10.1093/dnares/dsx042 |
| Angiosperms | *Rosa rugosa* | 952 | 10.1038/s41438-021-00594-z |
| Angiosperms | *Rosmarinus officinalis* | 2183 | MPGR database |
| Angiosperms | *Rubus chingii* | 879 | 10.1111/tpj.15394.聽 |
| Angiosperms | *Rubus corchorifolius* | 470 | GWH database |
| Angiosperms | *Rubus occidentalis* | 667 | 10.5524/100465 |
| Angiosperms | *Saccharum spontaneum* | 1905 | 10.1038/s41588-018-0237-2 |
| Angiosperms | *Saccharum spp* R570 | 153 | sugarcane database |
| Angiosperms | *Salix brachista* | 563 | 10.1038/s41467-019-13128-y |
| Angiosperms | *Salix purpurea* | 1543 | JGI database |
| Angiosperms | *Salvia bowleyana* | 604 | 10.1111/jipb.13085 |
| Angiosperms | *Salvia miltiorrhiza* | 676 | 10.1002/tpg2.20041 |
| Angiosperms | *Salvia splendens* | 1894 | 10.1093/gigascience/giy068 |
| Angiosperms | *Schrenkiella parvula* | 524 | 10.1104/pp.113.233551 |
| Angiosperms | *Sclerocarya birrea* | 469 | 10.5524/101057 |
| Angiosperms | *Scutellaria baicalensis* | 664 | 10.1016/j.molp.2019.04.002 |
| Angiosperms | *Scutellaria barbata* | 609 | 10.1016/j.gpb.2020.06.002 |
| Angiosperms | *Secale cereale* | 1402 | 10.1111/tpj.13436 |
| Angiosperms | *Senna tora* | 589 | 10.1038/s41467-020-19681-1 |
| Angiosperms | *Sesamum indicum* | 808 | 10.1186/gb-2014-15-2-r39 |
| Angiosperms | *Setaria italica* | 1209 | 10.1038/nbt.2196 |
| Angiosperms | *Setaria viridis* | 1556 | 10.1038/s41587-020-0681-2 |
| Angiosperms | *Simmondsia chinensis* | 648 | 10.1126/sciadv.aay3240 |
| Angiosperms | *Sinapis alba* | 1648 | 10.1371/journal.pone.0231002 |
| Angiosperms | *Siraitia grosvenorii* | 491 | 10.5524/100452 |
| Angiosperms | *Solanum aethiopicum* | 397 | 10.1093/gigascience/giz115 |
| Angiosperms | *Solanum chacoense* | 1099 | 10.1111/tpj.13857 |
| Angiosperms | *Solanum chilense* | 516 | 10.1534/g3.119.400529 |
| Angiosperms | *Solanum commersonii* | 645 | 10.5061/dryad.gf52g00 |
| Angiosperms | *Solanum lycopersicoides* | 751 | 10.1371/journal.pone.0242882 |
| Angiosperms | *Solanum lycopersicum* | 607 | 10.1101/767764 |
| Angiosperms | *Solanum melongena* | 545 | 10.1093/dnares/dsu027 |
| Angiosperms | *Solanum pennellii* | 688 | 10.1038/ng.3046 |
| Angiosperms | *Solanum pimpinellifolium* | 537 | 10.1093/dnares/dsaa029 |
| Angiosperms | *Solanum tuberosum* | 899 | 10.1093/gigascience/giaa100 |
| Angiosperms | *Sorghum bicolor* | 1187 | 10.1111/tpj.13781 |
| Angiosperms | *Spinacia oleracea* | 433 | 10.1002/tpg2.20101 |
| Angiosperms | *Spirodela polyrhiza* | 448 | 10.1038/ncomms4311 |
| Angiosperms | *Stanleya pinnata* | 731 | JGI database |
| Angiosperms | *Stevia rebaudiana* | 1339 | 10.1038/s41438-021-00565-4 |
| Angiosperms | *Striga asiatica* | 453 | 10.5061/dryad.53t3574 |
| Angiosperms | *Strobilanthes cusia* | 833 | 10.1111/tpj.14992 |
| Angiosperms | *Suaeda aralocaspica* | 399 | 10.5524/100646 |
| Angiosperms | *Syzygium oleosum* | 1283 | NCBI database |
| Angiosperms | *Taraxacum kok-saghyz* | 906 | 10.1038/s41598-019-38532-8 |
| Angiosperms | *Tarenaya hassleriana* | 906 | 10.1105/tpc.113.113480 |
| Angiosperms | *Tectona grandis* | 1358 | 10.5524/100550 |
| Angiosperms | *Theobroma cacao* | 911 | 10.1186/gb-2013-14-6-r53 |
| Angiosperms | *Thinopyrum elongatum* | 1288 | 10.1007/s00122-020-03591-3 |
| Angiosperms | *Thinopyrum intermedium* | 6540 | JGI database |
| Angiosperms | *Thlaspi arvense* | 706 | 10.1093/dnares/dsu045 |
| Angiosperms | *Toona sinensis* | 310 | 10.1111/1755-0998.13318 |
| Angiosperms | *Trichopus zeylanicus* | 499 | 10.1534/g3.119.400164 |
| Angiosperms | *Trifolium pratense* | 705 | 10.1038/srep17394 |
| Angiosperms | *Trifolium subterraneum* | 448 | 10.1038/srep30358 |
| Angiosperms | *Tripterygium wilfordii* | 1120 | 10.1038/s41467-020-14776-1 |
| Angiosperms | *Triticum aestivum* | 4679 | 10.1126/science.1251788 |
| Angiosperms | *Triticum dicoccoides* | 7153 | 10.1111/pbi.12940 |
| Angiosperms | *Triticum spelta* | 3476 | ensembl database |
| Angiosperms | *Triticum turgidum* | 6445 | 10.1038/s41588-019-0381-3 |
| Angiosperms | *Triticum urartu* | 914 | 10.1038/s41586-018-0108-0 |
| Angiosperms | *Urochloa fusca* | 1656 | JGI database |
| Angiosperms | *Utricularia reniformis* | 581 | 10.5281/zenodo.3268745 |
| Angiosperms | *Vaccinium corymbosum* | 487 | 10.1186/s13742-015-0046-9 |
| Angiosperms | *Vaccinium macrocarpon* | 606 | 10.3389/fpls.2021.633310 |
| Angiosperms | *Valeriana officinalis* | 755 | MPGR database |
| Angiosperms | *Vernicia fordii* | 893 | 10.1093/pcp/pcy117 |
| Angiosperms | *Vicia sativa* | 447 | 10.1101/2021.03.09.434686 |
| Angiosperms | *Vigna angularis* | 726 | 10.1073/pnas.1420949112 |
| Angiosperms | *Vigna radiata* | 1155 | 10.1038/ncomms6443 |
| Angiosperms | *Vigna subterranea* | 616 | 10.1093/gigascience/giy152 |
| Angiosperms | *Vigna unguiculata* | 1651 | 10.1111/tpj.14349 |
| Angiosperms | *Vitellaria paradoxa* | 428 | gdb database |
| Angiosperms | *Vitis arizonica* | 2304 | grapegenomics database |
| Angiosperms | *Vitis riparia* | 1276 | 10.1038/s41438-020-0316-2 |
| Angiosperms | *Vitis rotundifolia* | 1507 | 10.1093/g3journal/jkab033 |
| Angiosperms | *Vitis vinifera* | 1280 | 10.1038/nature06148 |
| Angiosperms | *Xanthoceras sorbifolium* | 640 | 10.5524/100606 |
| Angiosperms | *Zea mays* | 695 | 10.1105/tpc.16.00353 |
| Angiosperms | *Zingiber officinale* | 2845 | 10.1038/s41438-021-00627-7 |
| Angiosperms | *Ziziphus jujuba* | 1361 | 10.1038/ncomms6315 |
| Angiosperms | *Zostera marina* | 356 | 10.12688/f1000research.38156.1 |
| Angiosperms | *Zostera muelleri* | 490 | 10.4225/23/5ad41a6c77b60 |
| Angiosperms | *Zoysia japonica* | 215 | 10.1093/dnares/dsw006 |
| Angiosperms | *Zoysia matrella* | 355 | 10.1093/dnares/dsw006 |
| Angiosperms | *Zoysia pacifica* | 250 | 10.1093/dnares/dsw006 |

**Table S2. The information of 23 plant species with experimentally determined phosphorylation sites**

| **Species** | **The numbers of RLK+RLCK** | **The numbers of NRE segments** | **The ratio of NRE (%)** |
| --- | --- | --- | --- |
| *Arabidopsis thaliana* | 1150 | 1133 | 99 |
| *Brachypodium distachyon* | 1479 | 1421 | 96 |
| *Brassica napus* | 1951 | 1906 | 98 |
| *Chlamydomonas reinhardtii* | 4 | 4 | 100 |
| *Cicer arietinum* | 906 | 891 | 98 |
| *Citrus sinensis* | 1185 | 1160 | 98 |
| *Glycine max* | 2212 | 2174 | 98 |
| *Gossypium hirsutum* | 2565 | 2507 | 98 |
| *Hordeum vulgare* | 6503 | 6278 | 97 |
| *Jatropha curcas* | 830 | 820 | 99 |
| *Lotus japonicus* | 825 | 803 | 97 |
| *Medicago truncatula* | 1261 | 1230 | 98 |
| *Nicotiana tabacum* | 1633 | 1584 | 97 |
| *Oryza indica* | 941 | 888 | 94 |
| *Oryza sativa* | 1238 | 1205 | 97 |
| *Physcomitrella patens* | 1210 | 1198 | 99 |
| *Populus trichocarpa* | 1861 | 1780 | 96 |
| *Selaginella moellendorffii* | 782 | 771 | 99 |
| *Solanum lycopersicum* | 625 | 607 | 97 |
| *Solanum tuberosum* | 918 | 899 | 98 |
| *Triticum aestivum* | 4809 | 4679 | 97 |
| *Vitis vinifera* | 1300 | 1280 | 98 |
| *Zea mays* | 724 | 695 | 96 |

**Table S3. 31 motifs with experimentally determined phosphorylation sites in NRE segments**

| **Motif** | **Experimentally determined phosphorylation site** |
| --- | --- |
| CSSVYHHERAGSSQSGEEGSSGTVRKQSS | 2;3;5;12;13;15;20;21;23;28;29 |
| DDEEEETGCWVKLRFIGSCISSRSKVDSSISGT | 7;18;22;24;28;29;31 |
| GQSSNALSPSGSSYLTSSSEAAGILDSTVFEEREESECELE | 3;8;12;14;27;28;36 |
| IGMSNYRSDLDFSGNV | 8;13 |
| KEPLSINVATFEKPL | 5 |
| KLKZKFFKQNGGLLLQQQJSS | 20 |
| KVPASPLRVPPSPSRFSMSPKLNRJGSVH | 5;12;17;19;27 |
| MGCFSCFGSKK | 9 |
| MGNCLSSRIKAESPSSTGSSS | 6;7;13;15;16;17;19;20;21 |
| NIFTRKFKGPDVPSTVLKSAPDFCTVYVISKGKJ | 30 |
| PIRANGADSCTILSDSTIGPESPVKSGRNGMSLWLEGFRRSNVVSASGIP | 9;11;14;16;17;22;26;32;41;45;47 |
| PPPPPPPPPMMKSDPYGGQAFSWPHNPP | 13;16;22 |
| PSSASLSFGSSIAAYTGSAKT | 2;3;5;7;10;11;16;21 |
| QKEDLEKMKRQRDEEHSVAREQKSILESQ | 24;28 |
| QKRPQDWQKRNSFSSWLLPLHAGDSSFMSSKSS | 12;14;15;25;26;29;30;32;33 |
| RRRKKKLSGLNGGYVMPSPLGSSPRSDSSFTKTHSSAPLVG | 8;14;18;22;23;26;28;29;31;33;35;36 |
| SAVELLQKYKKERDELQVERDNALKEAEELRKKQAEASSSH | 9;40 |
| SIRGPVVTPTSSPELGTPFTATEAGTSSVSSSDPGTSPFFI | 1;8;10;11;12;17;20;22;26;27;28;30;31;32;36;37 |
| SQIPNVSKEIKEDRV | 1;7 |
| SQPKVLRLNLVGSPKKEPE | 13 |
| SSGGRSEKPKEEFGSGVQEAEKNKL | 1;6;15 |
| SSGWLPLPLYGNSH | 1;2;10;13 |
| SSSSKVSSASVPPTPRSE | 1;2;3;4;7;8;10;14;17 |
| SWPWKLTAFQ | 7 |
| SZFKASVLEAPDVENEEKSEVDB | 1;6;19 |
| TGSYASSLPSBLCR | 1;3;4;6;7;10 |
| TGYSSKLLABARYISQTMKLGALGLPAY | 1;5;28 |
| TKNGBLFSIWNFD | 1 |
| TSSEQKSDITDSCSQMILQLHDVYDPNKINVKIKIVSGSPC | 1;12 |
| VGPWKTGLSGQLQKAFVTGVP | 6;9;18 |
| ZELEKMKNQRDEVMEELQMALDQKSSLESQIAESDQMVKEL | 29;34 |
